# Quantifying phenology in the deciduous tree and phytophagous insect system: a methodological comparison

**DOI:** 10.1101/2025.01.30.635661

**Authors:** Lucy M. Morley, Sam J. Crofts, Ella F. Cole, Ben C. Sheldon

## Abstract

The extent to which phenological synchrony between trophic levels may be disrupted by environmental change has been a topic of increased focus in recent years. Phenological associations between deciduous trees, phytophagous insects and their consumers (e.g. passerine birds) have become one of the model systems for understanding this process. However, most existing research reports population-level associations rather than examining the smaller spatial scales at which these trophic interactions occur. Furthermore, a variety of methods have been used to measure phenology, particularly on producers and primary consumers, with little formal comparison. To investigate how different methods of measuring producer and primary consumer phenology influence our understanding of these biological relationships at the appropriate scale, we quantified phenological metrics for individual host trees and the phytophagous insects that depend on them in a deciduous woodland during spring 2023. We sampled 170 trees from six deciduous species in Wytham Woods, UK, deriving nine metrics of phenology from five distinct field methods: multispectral drone imaging (NDVI), hemispherical canopy photography, and bud-scoring observations to track tree phenology, as well as water traps and frass traps to monitor insect herbivore phenology. We assessed the reliability of these methods within both trophic levels and across tree species. We further evaluated the extent to which tree phenology metrics correlated with herbivore phenology, at the level of individual trees, and links to variation in subsequent herbivory rates across a subsample of 72 oak trees (*Quercus robur*). Our results illustrate how methodological choices can affect our ability to study the timing of trophic interactions and reveal finescale spatiotemporal variation in phenology across both trophic levels. We discuss the implications of these results for considering how the scale-dependence of trophic interactions may stabilise populations and shape broader-scale responses to environmental change.

## Introduction

Phenology, the timing of seasonal life history events, can be measured in many ways across different taxa and particular events. For interacting species, differential rates of phenological change due to the disruptive effects of climate change can result in phenological mismatch (Thackeray et al., 2016; Chmura et al., 2018; Cushing, 1969). Such divergence in event timing may disrupt ecosystem functioning and compromise both individual fitness and population-level survival (Samplonius et al., 2021; Visser & Gienapp, 2019). Hence, it is paramount that we can measure phenology with methods that are both accurate and at the appropriate scale.

Phenological trends and the assessment of mismatch are mostly quantified over long time periods and/or at broad spatial scales using population-level measures (e.g. Menzel & Fabian, 1999; Burgess et al., 2018; Charmantier et al., 2008). However, the interactions between organisms that determine the outcome of any phenological mismatch typically occur over small spatial and temporal scales, therefore the extent of mismatch may show fine-scale spatiotemporal variation depending on individual organisms’ responses to environmental cues and selective or plastic potential to maintain synchrony (van Asch et al., 2007; Kramer, 1995; Hinks et al., 2015; van Dongen et al., 1997). More simply, the fitness consequences of mismatch result from interactions between individuals, but this spatial scale-dependence of individual-level phenological synchrony has largely been neglected (Burgess et al., 2018; Hinks et al., 2015). Understanding synchrony at biologically relevant scales requires us to identify appropriate methods for quantifying phenological variation of individual organisms, both within and between species in a given system (Kharouba & Wolkovich, 2020).

Phytophagous invertebrate larvae and the deciduous trees on which they feed have become a popular model system for studying phenological synchrony in trophic plant-invertebrate interactions. Quantitative investigations into the potential for mismatch (asynchrony) in this system have been prompted by observations of differential phenological shifts and identification of underlying phenological cues. Both temperature and photoperiod determine tree leaf development (Vitasse & Basler, 2013; Vitasse et al., 2011), but invertebrate phenology appears more sensitive to temperature than plants owing to their ectothermic physiology (Forrest, 2016; Abarca & Spahn, 2021; Salis et al., 2016; Gutiérrez & Wilson, 2020). Leaf emergence dates not only vary between tree species, but amongst individuals of the same species (although individuals have been shown to be consistent between years), which can create a spatially and temporally varying selective environment for phytophagous insects whose fitness depends on hatching out in synchrony with their host tree’s budburst (Malyshev et al., 2022; van Dongen et al., 1997; Crawley & Akhteruzzaman, 1988; Wesołowski & Rowiński, 2006; Cole & Sheldon, 2017; though see Weir & Phillimore, 2024). Furthermore, many of these invertebrate species are generalists able to feed on multiple potential host tree species, so they must cope with both intra- and inter-specific variation in tree budburst (Tikkanen et al., 2000). Hence the timing of larval emergence is complex due to multiple environmental factors.

The emergence time of invertebrate larvae is under selection to coincide with the transient peak availability of their food, particularly where there is a trade-off between leaf size and the build-up of herbivory defence compounds (van Asch & Visser, 2007; Tikkanen & Julkunen-Tiitto, 2003). Winter moth larvae (Operophtera brumata (Geometridae)) have been a particular focus in the tri-trophic temperate woodland system due to their high abundance on pedunculate oak (Quercus robur), their role as a key food resource for passerine birds, and early work on this species as a model in population ecology (Shutt et al., 2019; Singer & Parmesan, 2010; Varley & Gradwell, 1958). Evidence suggests that, in some populations, synchronisation between oak and winter moth timing is becoming disrupted as their phenology advances to different extents with warmer springs (Visser & Holleman, 2001). However, selection may be acting fast enough in some winter moth populations to counter it (van Dis et al., 2023; van Dongen et al., 1997; van Asch et al., 2013). Some degree of toleration to both starvation and mature leaves revealed in recent experiments, and dispersal to other host trees (of the same or alternative species), may also buffer caterpillars against asynchronous hatching phenology (Weir, 2024; Weir & Phillimore, 2024).

A diverse array of methodological approaches has been used to collect data on the timing of phenological events in trees and phytophagous larvae. However systematic comparisons of these methods, how they agree, and how they capture variance among individuals across tree species are lacking. In this study, we explore the relationships among five field methods performed on the same individual trees to understand how the choice of method directly affects the values of timing metrics we extract from them, and how that may influence findings, interpretation and comparability of phenological data.

### Methods to quantify tree phenology

There are three main methods of studying tree phenology at varying spatial and temporal scales: (i) ground observations, (ii) aerial remote-sensing (e.g. satellites), and (iii) canopy-level remote-sensing (e.g. UAVs and PhenoCams; Polgar & Primack, 2013; Katal et al., 2022).

Ground-based observations of bud-to-leaf progression have historically served as a simple and practical way of recording individual tree phenology in the field, over long time periods or for reasonably large sample sizes (Crawley & Akhteruzzaman, 1988; Hunter & Lechowicz, 1992; van Dongen et al., 1997; Wesołowski & Rowiński, 2006; Hinks et al., 2015; Cole & Sheldon, 2017; Chamberlain & Wolkovich, 2021; Heinecke et al., 2024). Whether recording leaf emergence on a categorical scale to estimate budburst date or as percentage leaf cover, this method relies on repeated human observations at ground level.

Alternatively, imaging-based technologies offer more quantitative and, potentially, less subjective estimates of phenology. Multispectral sensors attached to satellites such as MODIS, Landsat, and Sentinel-2 generate remotely-sensed vegetation indices such as EVI (enhanced vegetation index) and NDVI (normalised difference vegetation index). From time series of images, we can extract green-up dates (e.g. when NDVI crosses 50% of its maximum or the date of maximum change in NDVI) and compare them between pixels (White et al., 2014; Zhang et al., 2003; Archibald & Scoles, 2007; Cole et al., 2015). However, satellite resolution (many metres) is too large to measure individual tree crown phenology, and their temporal resolution may also be insufficient. The recent increased availability of unmanned aerial vehicles (UAVs/drones) carrying RGB or multispectral sensors can generate high resolution (centimetre-level) time series image data at the intervals required to pick up temporal changes in vegetation indices (Fawcett et al., 2020; Berra et al., 2019; Park et al., 2019). UAV imagery also has the potential to scale up to measure crown NDVI of virtually all trees in the image with advancing crown segmentation and species detection algorithms (Weinstein et al., 2019; Nevalainen et al., 2017; Beloiu et al., 2023; Braga et al., 2020).

Digital cameras can also be used to collect repeat images from the ground or canopy-level mounts, known as “PhenoCams”, from which vegetation indices can similarly be extracted (Khare et al., 2022; Klosterman et al., 2014; Delpierre et al., 2020). Cameras fitted with hemispherical (fisheye) lenses capture images with a 180° field of view beneath the tree canopy (Chianucci, 2020; Smith & Ramsay, 2018). They are often used in forest structure research but may present an under-utilised method of monitoring tree phenology by tracking both greenness (green chromatic coordinate (GCC)) and canopy closure as leaves emerge, as software classifies pixels into sky vs canopy to calculate canopy attributes including percentage openness or leaf area index (LAI) (Chianucci & Macek, 2023; Atkins et al., 2020; Brown et al., 2020). This may present a lower cost option to monitor canopy phenology with lower data storage requirements than UAV imagery.

There is a need to understand how well simpler traditional methods relate to sophisticated new imaging-based technologies to ensure robust inferences are drawn from integrating different methods over temporal and spatial scales. For trees, some studies validating image-derived measures against ground-truthing observational data suggest there is very good agreement but have found offsets in the absolute dates of events (e.g. Klosterman et al., 2018; Berra et al., 2019; Soudani et al., 2008; Delpierre et al., 2020; Chianucci & Macek, 2023; Atkins et al., 2020).

### Methods to quantify primary consumer phenology

Several methods have been deployed to characterise the phenology of phytophagous insect larvae (henceforth caterpillars), depending on the goals of the study. Considerable focus on the winter moth, which is frequently the dominant species in these assemblages over the long term despite population cycles, has led to methods designed to assay their phenology in particular. Winter moths descend to the ground to pupate in soil, so one direct method of trapping them, and other descending species, is to intercept them with a water-filled tray on the ground. The half-fall (median) date, when 50% of (fifth final instar) larvae have been intercepted between tree and ground, serves as a standard metric of their phenology (van Noordwijk et al., 1995; Charmantier et al., 2008; Hinks et al., 2015). This metric is strongly correlated with spring temperatures and has advanced at a similar rate to the mean lay date of great tits (*Parus major*), which rely on winter moths as an important food source for their nestlings at our study site Wytham Woods (van Noordwijk et al., 1995; Charmantier et al., 2008). In addition to winter moth larvae, this method also captures other taxa incidentally when these are displaced from tree canopies. Branch-beating similarly collects individual larvae falling from trees but involves actively hitting the branches to make larvae fall (Shutt et al., 2019). This requires low hanging accessible branches, which may not always be present, and may not capture invertebrate phenology that is representative of the whole tree’s canopy due to microclimate and physical isolation influencing the within-canopy distribution of larvae. Alternatively, biomass of the whole caterpillar community can be estimated indirectly from frass traps – receptacles collecting caterpillar faecal pellets. This method has been widely used to infer the abundance peak as a measure of caterpillar timing and densities (Tinbergen & Dietz, 1994; Zandt, 1994; Seress et al., 2018; Burgess et al., 2018; Nadolski et al., 2021; Veen et al., 2010).

Both frass and water traps could be used to extract half-fall and peak dates but have rarely been explicitly compared. For example, a small sample of paired frass and water traps show strong positive correlation in peak dates (*F*_1,11_=82.0, p<0.0001, *R*^2^=0.88; Mallord et al., 2017). Hinks et al (2015) found positive associations between both peak frass and winter moth half-fall dates and the budburst date of their host oaks (half-fall: *F*_1,61_=73.7, p<0.001, *R*^2^=0.67; peak frass: *F*_1,75_=26.2, p<0.001, *R*^2^=0.57), but samples were taken from one area within Wytham Woods rather than across multiple sections of the site which vary in habitat and microclimate.

To our knowledge, no study has explored the relationships between bud scoring, hemispherical photography and multispectral imaging to measure tree phenology whilst simultaneously measuring caterpillar phenology with both frass and water traps at the individual tree level. Therefore this is the first direct comparison of these multiple methods for quantifying phenology across both trophic levels performed on the same individual trees, shedding light on the implications of methodological choices for understanding biological relationships of trophic interactions. Our aims were to:

1) Quantify phenological metrics for individual trees in conjunction with their associated phytophagous invertebrate communities using multiple field methods.
2) Examine the relationships between phenological metrics derived from different field methods across six deciduous tree species by testing:
  a) the correlation of metrics to assess how well these methods capture tree and caterpillar phenology across tree species (i.e. methodological comparisons within trophic levels),
  b) the extent to which tree phenology (measured in different ways) can predict the timing of caterpillar phenology on individual oak trees (i.e. biological relationships across trophic levels).
3) Examine the relationships between phenological metrics and primary consumer productivity by testing the extent to which tree and caterpillar phenology (measured in different ways) can predict herbivory rates of individual oak trees as the consequence of trophic synchrony.

## Methods

### Study site and tree selection

We conducted fieldwork at Wytham Woods (51°46N, 1°20W, National Grid Reference SP4608), a 385ha temperate deciduous woodland in Oxfordshire, UK, in March-June 2023. We selected 170 mature trees of six common species: ash (*Fraxinus excelsior*; n=23), beech (*Fagus sylvatica*, n=5), silver birch (*Betula pendula*, n=17), hazel (*Corylus avellana*, n=30), pedunculate oak (*Quercus robur*, n=77), and sycamore (*Acer pseudoplatanus*, n=18; see **Figure 1** for a map). To capture greater biological complexity, we included additional tree species beyond the classic simplified oak – winter moth pairing, because although winter moths are particularly numerous on oaks, they do feed on other deciduous trees, and other phytophagous larvae also consume oak leaves (Tikkanen et al., 2000; Shutt et al., 2019; Weir, 2024; Macphie et al., 2024). Our focal trees were distributed based on a 100m x 100m grid of Ordnance Survey reference posts in two plots within Wytham: Marley Wood (southeast) and Great Wood (northwest). In Marley, one tree per species (if present) was selected around each of 22 grid posts, totalling 78 trees. In Great Wood, we similarly chose one tree per non-oak species (if present) around 12 posts, but we chose three oaks per post to investigate phenological variability at even finer spatial scales in other analyses, so in this part of the site we selected a total of 77 trees. To eliminate phenological bias, we selected these trees in early March 2023 before any leaf development. Additionally, we sampled 25 oaks in the centre of the woods (ForestGEO plot) not based around the grid posts. These trees have been previously studied, enabling the estimation of between-year phenological relationships in other analyses. We used Emlid Reach RS2 receivers (which correct position drift error from GPS satellites) to obtain high precision longitude and latitude coordinates for each tree.

**Figure 1.**
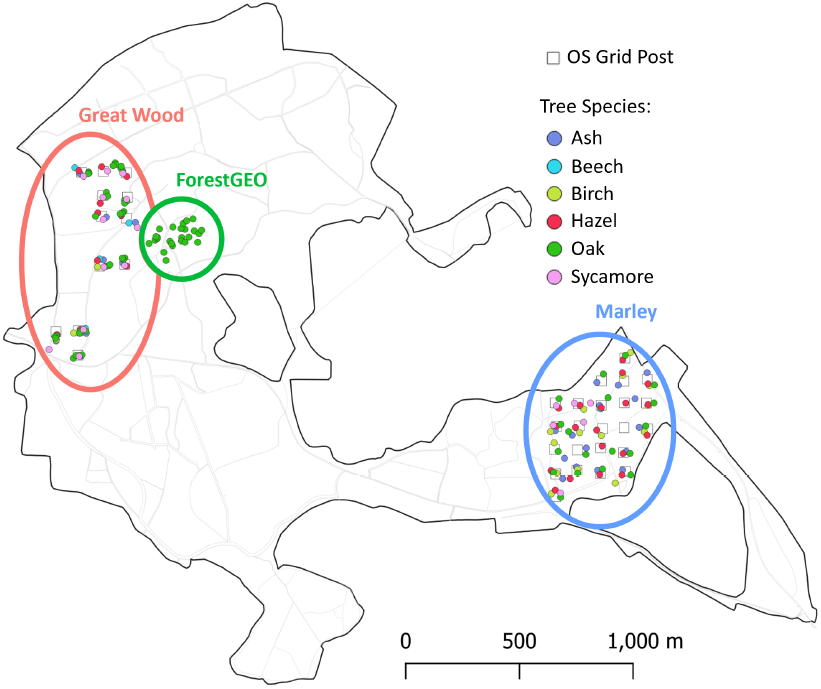
Map of Wytham Woods showing the locations of 170 trees sampled for tree phenology and their associated caterpillar community phenology, distributed within three areas of the site referred to as Great Wood (red), ForestGEO (green), and Marley (blue). Marker colour denotes tree species and black squares are Ordnance Survey 100m-grid posts. Gaps of >100m occur in Great Wood and Marley where paths or thick understorey prevented selection of suitable trees. Ash n=23, beech n=5, birch n=17, hazel n=30, oak n=77, and sycamore n=18.

To monitor phenology of individual trees and their caterpillar communities, we used five field methods for which we collected data at approximately three-day intervals throughout the 2023 spring season (**Figure 2**; **SI Table 1**).

**Figure 2.**
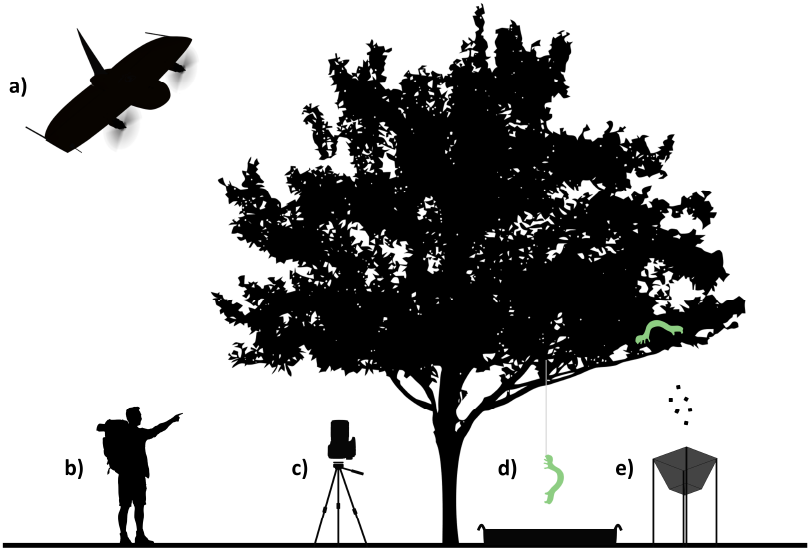
Schematic of field methods performed on each tree: a) multispectral drone imaging, b) bud scoring observations, c) hemispherical canopy photography, d) caterpillar water traps, and e) frass traps.

### Multispectral imaging with drones

To capture multispectral image data of the whole tree canopy of Wytham Woods, we flew WingtraOne Gen II drones carrying multispectral MicaSense RedEdge-P cameras at 100m above ground level (ground sampling distance = 3.4cm/pixel). We split the site into 12 flight areas which we surveyed every 3-4 days (depending on wind speed and precipitation being below operating limits) between 21^st^ March and 9^th^ June 2023, yielding a time series of 21-23 orthomosaics per flight area. To maximise the precision of stitching quality of our image orthomosaics, we applied post-processing kinematics (PPK) to the image location data using GPS-corrections from an Emlid Reach RS2 base station placed at the centre of our study site. Its fixed position was known to centimetre-level, so it greatly improved our geotagging accuracy by negating GPS drift. We then stitched the images with panchromatic sharpening using PIX4Dfields to construct NDVI orthomosaics for each flight (see Wingtra, 2025 for details on drone data acquisition and processing). We manually delineated shapefiles of the focal tree crowns and passed the series of orthomosaics through a pipeline in ArcGIS Pro to extract mean NDVI for the core 75% of each crown. We chose 75% of the crown area to minimise edge effects of branches shifting between flights, e.g. due to wind conditions. No NDVI data was acquired for hazel trees as their crowns were below the canopy and therefore did not appear in the drone images.

### Bud scoring

To monitor bud progression, we used photographic keys (developed by Hinks et al. (2015) and Cole & Sheldon (2017)) to assign a bud development stage to each tree (the average score of 12 crown sections; four horizontal quarters of the lower, middle, and upper strata). Scores corresponded to which stage the crown section showed the dominant characteristics of, ranging from 1 (no or minimal bud swelling), to 5 (for non-oaks) or 7 (for oaks) denoting full leaf (photographic references available in **SI Figure 1**). Stage 3 (for non-oaks) and stage 4 (for oaks) correspond to budburst, i.e. when leaves first start emerging as the buds open. Primarily dead sections were recorded as NA. To mitigate observation bias, observers were trained by the same ‘expert’ in early March and rotated around sampling areas. Twenty rounds of observations were conducted within the sampling period from 25^th^ March to 25^th^ May 2023.

### Hemispherical canopy photography

To obtain 180° raw-format images of tree canopies, we used Canon EF 8-15mm hemispherical lenses attached to Canon EOS 600D DSLR cameras. Cameras were mounted facing upwards on tripods set to 60cm, spirit-levelled, and with the top oriented north above a ground marker stake to ensure consistent positioning beneath the same portion of canopy every visit. We used aperture priority mode, with f=5.0, ISO=100 to minimise sensor noise, evaluative metering and 10mm focal length for full-frame images. These have greater resolution than circular images, remove some non-focal canopy from the edges of frame, and correlate better with field sampling estimates of leaf area (Macfarlane et al., 2007). Images were taken at +1/3, 0 and -1/3 exposure compensation (EV) to account for variable light conditions and brightness histograms were checked in-field to ensure no extreme skew (Beckschäfer et al., 2013). Ideally, images were captured under overcast skies or without direct sun in frame to minimise glare and over-exposure. Logistically, this was not always possible. To control for weather conditions and inform outlier identification, we recorded weather conditions as sun, overcast, or a mixture of mainly sun/some cloud or mainly cloud/some sun. Heavy rain days meant 16-22 rounds were conducted within the sampling period (25^th^ March to 8^th^ June) depending on area.

All images were processed with a gamma-adjusted contrast stretch of the raw file’s blue channel in bRaw to standardise exposure variation to reduce bias in canopy attribute estimates (Chianucci, 2022). The resulting JPEGs were processed in hemispheR to compute canopy attributes, including diffuse non-interceptance (% canopy openness), and actual and effective leaf area index (LAI), with parameters set to 7 zenith rings, 8 azimuth sections, and zonal thresholding to accommodate heterogenous sky conditions (Chianucci & Macek, 2023). Slightly under-exposed -1/3EV images were used for analysis (Brusa & Bunker, 2014; Macfarlane et al., 2014). We used effective LAI for analysis because it accounts for three-dimensional clumping (Chianucci & Macek, 2023). We identified and removed extreme outliers through visual inspection (n=115) arising from incorrect camera settings, sun glare, or binarization errors to leave a total of 2,839 images.

### Caterpillar water traps

Individual larvae were collected via water traps placed under the tree canopy. Traps consisted of a 100cm x 55cm (0.55m^2^) plastic tray filled with water to approximately 10cm in depth to submerge descending larvae. Traps had a wire mesh lid to prevent birds from feeding on the larvae. For each sampling round, we sieved through the contents to remove the dead larvae and recorded their abundance under the following categories: non-final instar winter moth, final instar winter moth, green oak tortrix (*Tortrix viridana*), other green Lepidoptera, and red/brown Lepidoptera, and Symphyta (sawfly larvae). To minimise misidentification, observers were trained in distinguishing these groups and equipped with photographic references in the field (available in **SI Figure 2**). We visited traps fourteen times within the sampling period (3^rd^ May to 20^th^ June) and stopped when the last two visits yielded only one or no caterpillars.

### Frass traps

To collect caterpillar excrement, we positioned frass traps >1m away from water traps under the tree canopy to avoid interference. Traps comprised of muslin cloth attached to a 50cm x 50cm (0.25m^2^) wire frame on bamboo canes 1m off the ground, with a stone to weigh down the cloth allowing matter to accumulate at the bottom. During each visit, we carefully scraped the frass and leaf litter into paper envelopes. We collected frass under each tree eleven times within the sampling period between 9^th^ May and 15^th^ June 2023. At the end of the season, samples were oven-dried at 60°C for 24hrs to remove water content. Dried frass was then sorted from litter with 1mm and 0.5mm sieves and weighed to the nearest 0.01g.

### Herbivory

In July 2023, after the main caterpillar spring phenological distribution, we returned to 72 of the 77 oak trees we had monitored for phenology to gather additional data on herbivory levels (five trees were not sampled due to inaccessible canopies). We used long-handled shears or an arborist’s slingshot to retrieve a bunch of 10 leaves per tree. The relative canopy level (lower, middle, upper) the branch came from was recorded. For each leaf, we measured total leaf area (cm^2^), actual consumed leaf area (cm^2^), and percentage consumed using the LeafByte mobile app on photos taken with an iPhone 12 Pro Max (Getman-Pickering et al., 2020).

### Statistical analyses

#### Calculating phenological metrics

To make cross-method and cross-species phenological comparisons, we calculated the dates of a) the half-date (median date) of each phenological trajectory and b) peaks in caterpillar abundance over the season (**Figure 3**). To estimate half-dates, we normalised the data to between 0 and 1 for each method per tree: 0 represented the minimum bud score of 1, the minimum effective LAI and mean NDVI values, and zero caterpillars and frass. Values of 1 corresponded to bud stage 5 (non-oaks) or 7 (oaks), the maximum effective LAI and mean NDVI values, and the cumulative total caterpillar and frass quantities collected by the end of the season. Half-dates for each method are referred to as follows: budburst for bud scoring, half-leaf for hemispherical photography, half-NDVI for multispectral drone imaging, and half-fall for caterpillar (repeated for all total caterpillars and final instar winter moths) and frass collection i.e. when 50% of the total amount of caterpillars or frass has fallen (van Noordwijk et al., 1995; **Figure 3a**). To generate a sigmoidal curve for each tree’s trajectory, we fitted Beta-family (logit link) generalized additive models (GAMs) in brms, with TreeID as a smoothing spline (factor-smooth interactions were used for bud scores, leaf area index, NDVI, frass, and total caterpillars, but individual smooths for final instar winter moths to avoid overfitting), and extracted the day of the year at which each curve passed through y=0.5 (half-date) in R v4.4.1 (Douma & Weedon, 2019; Bürkner, 2017; R Core Team, 2024). Bayesian methods enable incorporating uncertainty and extraction of lower and upper 95% credible intervals around the half-date estimates (**SI Table 2**), and we extracted the gradient at y=0.5. All models were fitted to each method and species separately given there were species-specific differences, and bud scoring models also included a random intercept for observer identity.

**Figure 3.**
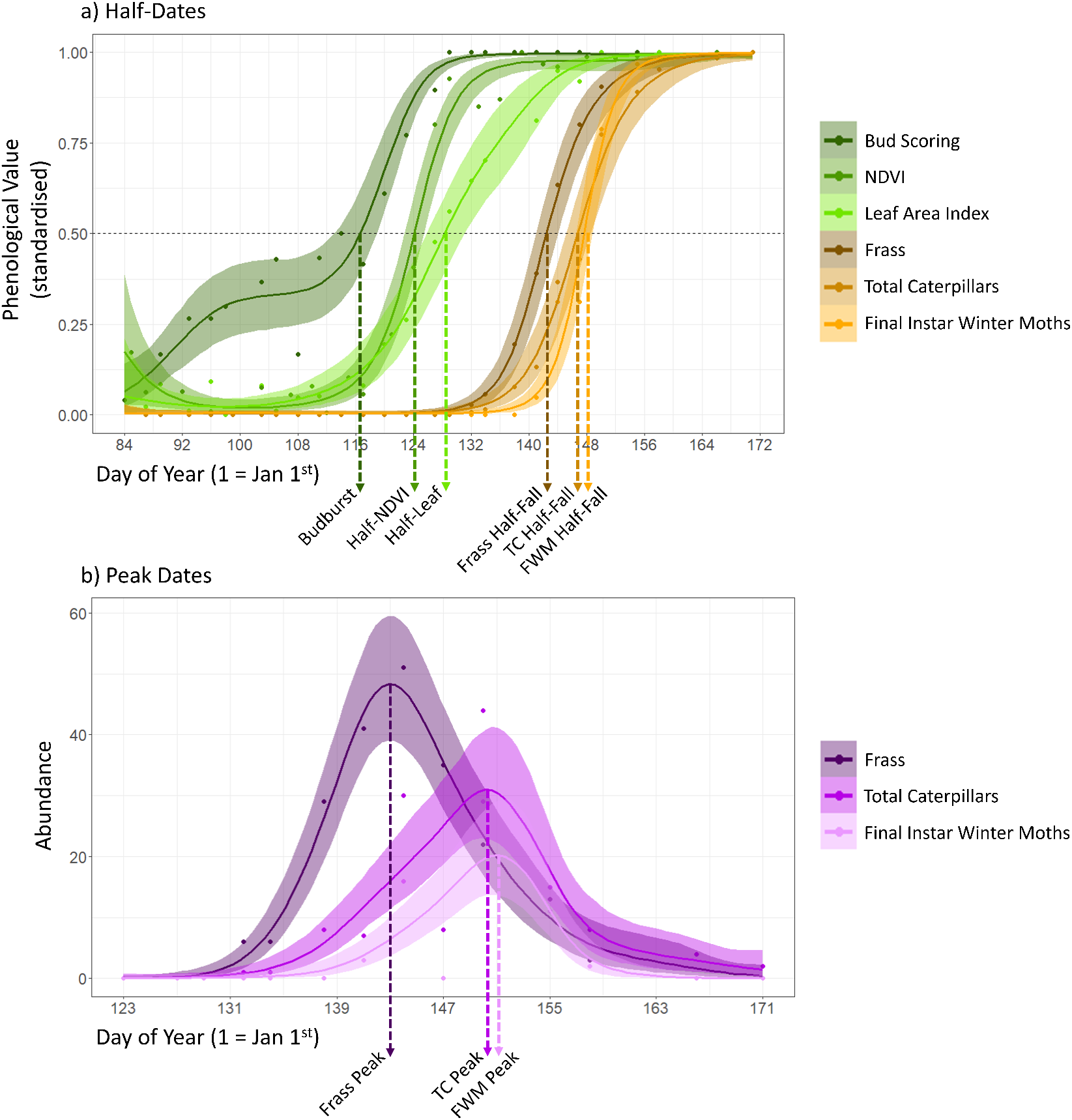
Example of a) half-date and b) peak date extraction for a single oak tree in this study. Note in b), frass mass is scaled in centigrams to enable fitting Poisson models to be comparable to caterpillar abundance data. TC = total caterpillars, FWM = final instar winter moth caterpillars.

We used a similar model framework to extract peak dates for caterpillar phenology. We fitted Poisson-family GAMs in brms to the raw counts of total caterpillars, final instar winter moths, and frass quantities over the sampling period smoothed by individual tree identity (as a factor-smooth in all cases) within each species. Frass mass was multiplied by 100 to transform it from gram to centigram scale so it was suitable to be fitted to the same model specification. For each tree, we extracted the peak height (maximum abundance; and its lower and upper 95% credible intervals) and the day of year on which the peak occurred. In total, there are nine phenological metrics derived from five field methods in this comparison study. It should be noted that values for many other aspects of phenology could be extracted, but we chose half-dates and peaks for simplicity and comparability with previous work as they are widely used standard measures of phenology (White et al., 2014; Briedis et al., 2024; Macphie et al., 2023; Fawcett et al., 2020).

#### Repeatability of field measures

It was possible to assess repeatability for four of the five field measures. Nine trees (five oaks, two birches, one ash, and one beech) were covered by two drone flight areas, allowing us to compare two independent sets of NDVI measures for these trees. We also used two identical camera setups to take hemispherical canopy photos in quick succession for the 25 ForestGEO oaks at the start of spring on 28^th^ March 2023 (i.e. in the early stages of bud emergence). Previous work has established that budburst dates quantified by this method (at this site) have moderate to high repeatabilities (Cole & Sheldon, 2017). Therefore, we did not test the inter-observer repeatability of bud scores here, but instead controlled for their potential variation by including observer as a random effect in the budburst extraction models. For the caterpillar measures, two oaks had two frass and two water traps throughout the sampling period. We assessed repeatability in terms of absolute agreement using the intraclass correlation coefficient (Gamer et al., 2019). Only one set of data was chosen to remain in the main analyses to avoid pseudoreplication.

#### Comparing phenological metrics

We summarised species-level variation in phenological metric values then constructed (Spearman) correlation matrices to assess within and between trophic level agreement in dates at the individual tree level using the corrplot package (Wei & Simko, 2024). Then we used brms models to explore: a) the effect of tree metric on the six caterpillar phenology metrics, for oaks and non-oaks (Gaussian mixed models), and b) the effect of tree or caterpillar phenology metrics on the mean percentage leaf area consumed i.e. herbivory (Beta mixed models; Bürkner, 2017). Models were compared based on effect strength, proportion of variation explained (conditional Bayesian *R*^2^) using the performance package (Lüdecke et al., 2021), and ELPD-LOO (expected log pointwise predictive density leave-one-out cross-validation) to assess robustness. All models included the fixed effect of sampling area to control for broad-scale spatial differences and sampling strategy. Models were checked for residual spatial autocorrelation using Moran’s Index.

## Results

### Repeatability

The raw data obtained from field methods were all highly repeatable. Effective LAI, measured using two identical camera set ups that photographed the 25 ForestGEO oaks twice on the initial sampling round, yielded a repeatability of 0.85 (p<0.001; n=25 trees). Mean NDVI values of the nine tree crowns, measured from overlapping drone flights across the season, exhibited very high repeatability, with all trees showing R=0.89-0.99 and p<0.001, except for the ash (R=0.84, p<0.01; n=10-14 pairs of flights per tree). For caterpillar measures, the repeatability of the raw amount of frass collected by two traps under two oak trees across the season was also extremely high, with R=0.97 and R=0.92 for each tree (p<0.001 and p<0.05 respectively; n=11 timepoints for both trees). Similarly, the numbers of total caterpillars and final instar winter moth larvae collected in water traps each round were highly repeatable for the two trees (total caterpillars: R=0.91 and R=0.97, both p<0.001; winter moths: slightly lower at R=0.89 and R=0.86, both p<0.001; n=14 timepoints for both trees). See **SI Figure 3** for visualisation.

### Species-level variation in phenological metric dates

The variation in date values from the nine phenological metrics for each tree species are illustrated in **Figure 4**. Means and standard deviations of the estimated dates, and the mean and standard deviation for the widths of credible intervals around estimated half-dates, are presented in **SI Table 2**. Mean budburst dates across species exhibited a substantial spread of 31.9 days, whereas half-NDVI and half-leaf days were more confined across 12.06 and 8.40 days respectively. Caterpillar metrics demonstrated even narrower spreads between species means: 2.50 days for frass half-fall and 3.54 for peak frass, 4.54 days for half-fall and 7.28 for peak of total caterpillars, and 5.13 for half-fall and 5.65 for the peak of final instar winter moth caterpillars. Hence, despite variability in budburst date, these data show markedly reduced temporal variability in consumer phenology for the same trees.

**Figure 4.**
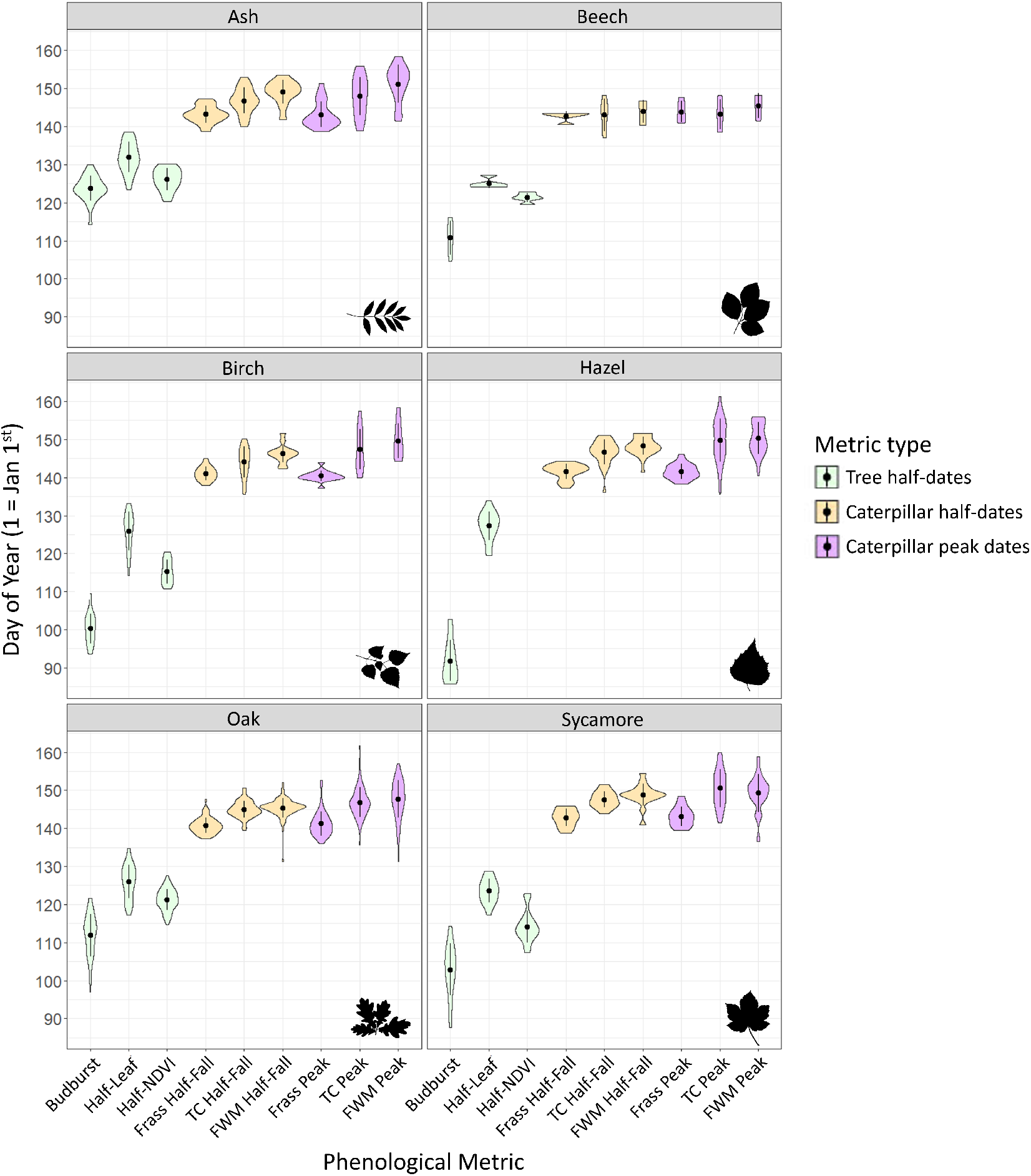
Distribution of day values (1 = 1st January) for nine phenological metrics for each tree species across 170 trees sampled in Wytham Woods. Green denotes tree half-date metrics (budburst, half-leaf, and half-NDVI day), whilst orange and purple are caterpillar half-dates and peak metrics respectively (frass, total caterpillars fallen (TC), and final instar winter moths (FWM)). Black circles and lines are means ± one standard deviation. Mean, standard deviation, and sample size values can be found in **SI Table 2**

The order in which species reach their half-dates varies across metrics within and between trophic levels (**Figure 4**; **SI Table 2**). However, within tree species, there was a consistent trend of budburst occurring earliest, followed by half-NDVI, and then half-leaf for tree-level metrics. At caterpillar level, the temporal spread was narrower, but again the pattern was consistent across species: frass occurred first, followed by total caterpillar, and then final instar winter moths, for both half-fall and peak dates (**Figure 4**). Tree species identity explained substantial proportions of variation in all nine metrics, though the proportion varied. Including tree species as a fixed effect (in addition to sampling area) increased the amount of explained variance (Bayesian *R*^2^) by 73.1% for budburst, 19.9% for half-leaf, and 58.8% for half-NDVI compared to models only controlling for sampling area. For caterpillar metrics, tree species explained 31.0% of the variance in frass half-fall, 22.3% in total caterpillar half-fall, and 33.3% in final instar winter moth half-fall. Peak date variances were lower, with *R*^2^ values of 18.6% for frass peak, 14.7% for total caterpillar peak, and 16.3% for final instar winter moth peak.

There was also variation in the mean width of credible intervals around half-date estimates among metrics and species (**SI Table 2**). Budburst had the biggest difference in CI width between species (hazel mean=11.52, ash mean=5.11), whilst half-leaf and half-NDVI were similarly wide within species and did not differ as much between species. For caterpillar metrics, frass half-fall had relatively narrow CIs across all species, ranging from 1.68 days for oak to 3.96 days for ash, while final instar winter moth half-fall CIs were much wider, particularly in ash (mean=14.18 days; **SI Table 2**).

### Methodological comparisons within trophic levels

For the tree phenology methodologies, metric type explained a substantial proportion of variation in estimated dates across all species. For oaks, metric type accounted for 76.2% of variation, calculated as the difference in conditional *R*^2^ between models controlling for tree identity (random effect) and sampling area, with and without the inclusion of metric type. In other species, this value was 55.6% for ash, 77.9% for beech, 86.9% for birch, 93.2% for hazel, and 78.6% for sycamore. Among caterpillar metrics, the amount of variation attributed to metric type was consistently lower than that observed for tree level. For oaks, metric type explained 46.0% variation in caterpillar metrics, 39.2% for ash, 3.6% for beech, 50.9% for birch, 62.4% for hazel, and 45.2% for sycamore.

In oak trees, there were significant positive correlations between most combinations of metrics apart from those involving half-leaf day (**Figure 5d**). Among the tree phenol-ogy metrics, oak budburst and half-NDVI dates showed the strongest correlation of any species (BB–NDVI Spearman’s r=0.84, p<0.001; n=76). Within caterpillar metrics, the pairs of half-fall and peaks for all methods (frass, total caterpillar, and final instar winter moth) were strongly correlated, which is expected as they are comparing statistical estimates derived from the same field data. Frass half-fall and total caterpillar half-fall showed the strongest cross-field method correlation amongst the combinations in oaks (FHF–TCHF r=0.70, p<0.001; n=77), followed by a slightly weaker relationship frass half-fall and final instar winter moth half-fall (FHF–FWMHF r=0.66, p<0.001; n=76; **Figure 5d**).

**Figure 5.**
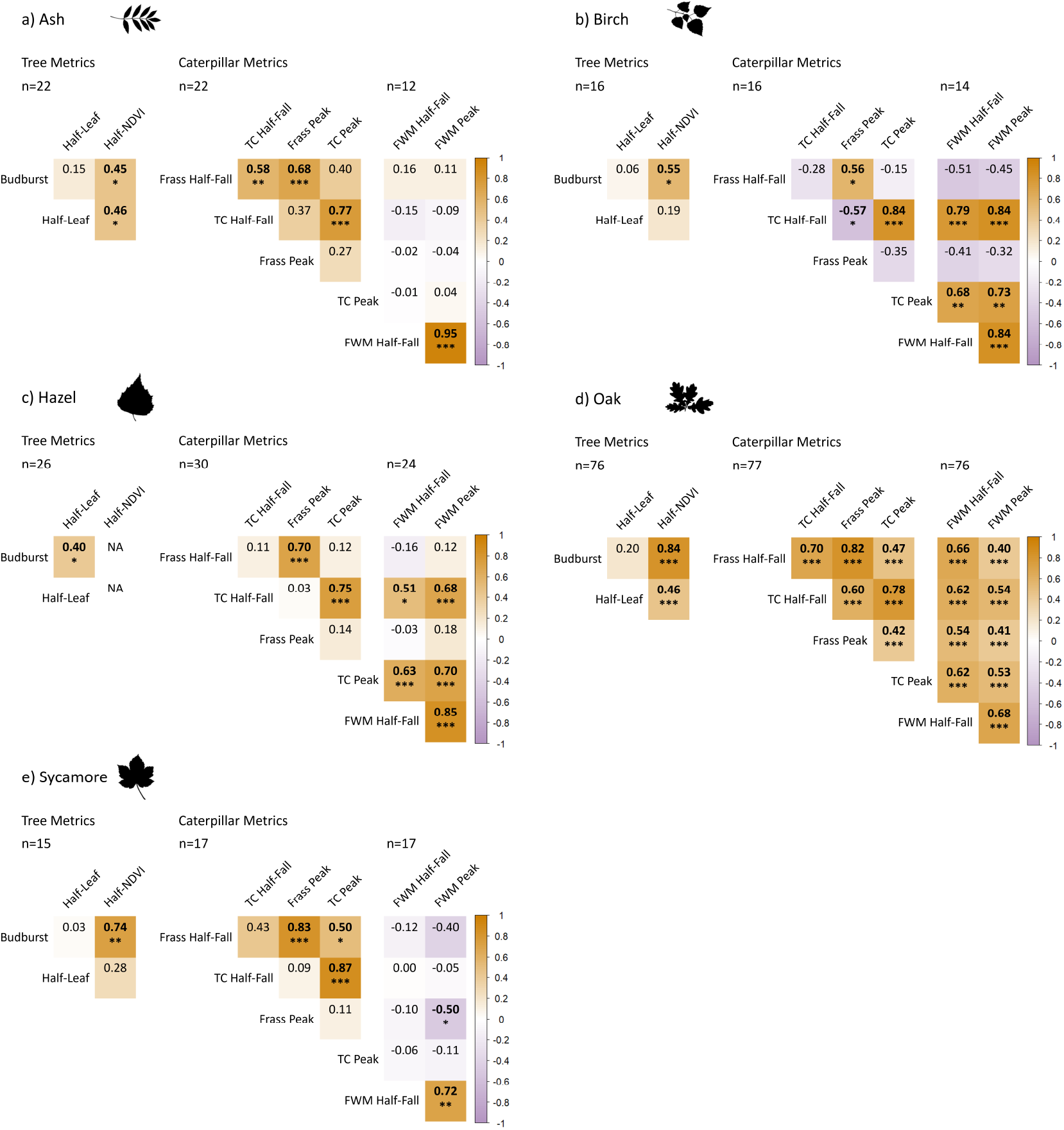
Correlation matrices between the nine phenological metrics for five tree species for within trophic level comparisons. Values are Spearman correlation coefficients; those in bold are significant at the p<0.05 threshold and asterisks denote significance thresholds: * = p<0.05, ** = p<0.01, *** = p<0.001. Differing sample sizes occur as few trees did not collect any caterpillars and several did not collect any final instar winter moths. For beech, results are less reliable due to limited sample size (n=5) so these are presented in **SI Figure 4**

For the other tree species, there were fewer significant correlations between metrics within trophic levels (**Figure 5a-c, e**), though sample sizes are lower so the estimates should be treated with caution. However, moderate correlations between budburst and half-NDVI (BB–NDVI) were observed in ash (r=0.45, p<0.05; n=22), birch (r=0.55, p<0.05; n=16) and sycamore (r=0.74, p<0.01; n=15). Budburst was only correlated with half-leaf in hazel (BB–HL r=0.40, p<0.05; n=26). At the caterpillar level, aside from the half-fall and peak pairings for each method, frass half-fall was correlated with total caterpillar half-fall in ash (FHF–TCHF r=0.58, p<0.01; n=22) and total caterpillar peak in sycamore (FHF–TCP r=0.50, p<0.05; n=17). Frass peak and total caterpillar half-fall were negatively correlated in birch (FP–TCHF r=-0.57, p<0.05; n=16), as was frass peak and final instar winter moth peak in sycamore (FP–FWMP r=-0.50, p<0.05; n=17). Total caterpillar half-fall and peak dates were positively correlated with final instar winter moth half-fall and peak dates in birch and hazel, although the relationship was stronger in birch (**Figure 5b & c**).

The significant correlations between total caterpillar metrics and final instar winter moth metrics in oak, birch and hazel reflect a part-whole relationship, as winter moths represent a subset of total caterpillar data (**Figure 5d, b & c**). The correlations for oak trees may be more precise than those for non-oaks due to the largest sample size (n=77) and higher caterpillar abundance influencing statistical power. The small sample size of beech trees (n=5) studied here precludes the estimation of correlations between phenological metrics with any precision. They are presented, for the sake of completeness, in **SI Figure 4**.

### Relationships between individual tree and caterpillar phenology

Oak budburst and half-NDVI dates showed moderate significant positive correlations with all caterpillar metrics (though half-NDVI was always slightly weaker), indicating later tree phenology is associated with later caterpillar phenology (**Figure 6d**). Meanwhile half-leaf day showed no significant correlations with caterpillar phenology. In other tree species, most combinations of tree and caterpillar metrics showed non-significant correlations (**Figure 6a-c, e**), except for a negative relationship between half-NDVI and final instar winter moth peak in ash (NDVI–FWMP r=-0.62, p<0.05; n=11), half-leaf and frass peak in hazel (HL–FP r=-0.43, p<0.05; n=26) and birch (HL–FP r=-0.54, p<0.05; n=15), and half-leaf and frass half-fall also in birch (HL–FHF r=-0.59, p<0.05; n=15). However, birch trees also showed significant positive correlations between half-leaf and total caterpillar half-fall (HL–TCHF r=0.72, p<0.01; n=15), half-leaf and final instar winter moth peak (HL–FWMP r=0.59, p<0.05; n=13), and half-NDVI with final instar winter moth half-fall (NDVI–FWMHF r=0.56, p<0.05; n=13; **Figure 6b**).

**Figure 6.**
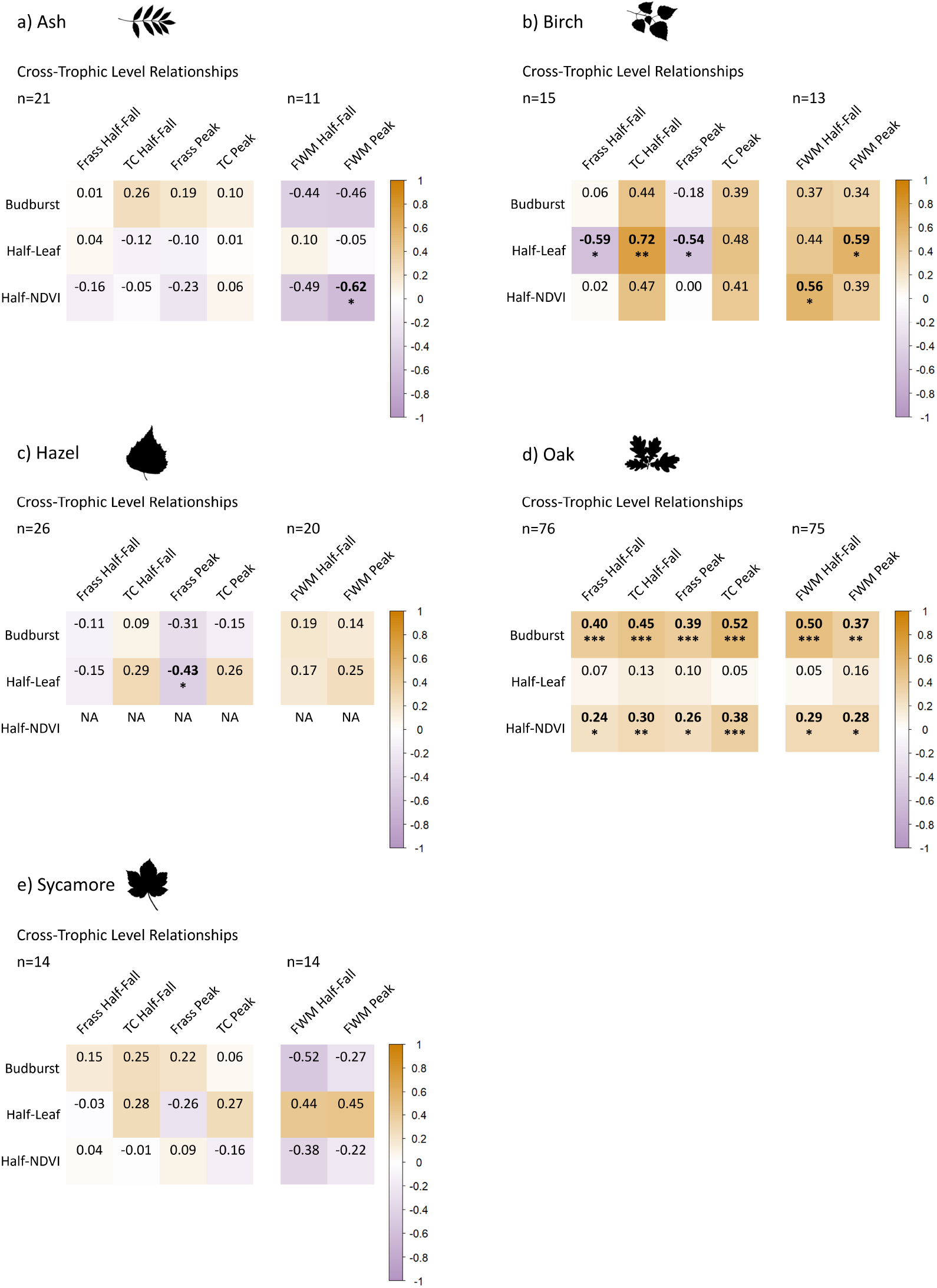
Correlation matrices between the nine phenological metrics for five tree species for between trophic level comparisons. Values are Spearman correlation coefficients; those in bold are significant at the p<0.05 threshold and asterisks denote significance thresholds: * = p<0.05, ** = p<0.01, *** = p<0.001. Differing sample sizes occur as few trees did not collect any caterpillars and several did not collect any final instar winter moths. For beech, results are less reliable due to limited sample size (n=5) so these are presented in **SI Figure 4**

### Spatial effects in predicting caterpillar phenology of host oaks

For oak trees, where we had measured a large sample of trees, we further tested the extent to which associations between tree and caterpillar phenology were consistent across space. Models controlling for the effect of sampling area revealed that both budburst and half-NDVI dates had significant positive main effects on all six caterpillar metrics, although half-NDVI had a stronger effect approximately twice the magnitude of the budburst effect (**Table 1**; **SI Figure 5**). Half-leaf consistently had the weakest effect and was only significant for total caterpillar (both half-fall and peak), final instar winter moth peak, and was marginally significant for their half-fall but had no significant effect on frass metrics.

**Table 1.** Outputs of Bayesian Gaussian models for the effect of tree phenology metrics on caterpillar phenology metrics for 77 oak trees sampled in Wytham Woods. All models included one of three tree phenology metrics and sampling area as fixed effects. Intercept is taken as area = Great Wood. 95%CIs are 95% credible intervals around the estimate and effects in bold are significant where CIs do not overlap zero. *R*^2^ is the amount of variation explained by the model. Δelpd is the difference in ELPD-LOO between the best fitting model (set to 0) and the remaining two models for each set of three tree metrics. HF = half-fall, TC = total caterpillars, FWM = final instar winter moth caterpillars.

|  |  | Estimate (95%CI) | R<sup>2</sup> (%) | Δelpd |  |  | Estimate (95%CI) | R<sup>2</sup> (%) | Δelpd |
| --- | --- | --- | --- | --- | --- | --- | --- | --- | --- |
| *Frass Half-Fall ~ Budburst* |  |  |  |  | *Frass Peak ~ Budburst* |  |  |  |  |
| n=77 | Intercept | **126.33 (119.48, 133.22)** | 47.7 | 0.0 | n=77 | Intercept | **118.75 (106.20, 131.66)** | 36.3 | -0.7 |
|  | Budburst | **0.12 (0.06, 0.18)** |  |  |  | Budburst | **0.19 (0.08, 0.30)** |  |  |
|  | FG | **2.32 (1.55, 3.12)** |  |  |  | FG | **3.36 (1.99, 4.76)** |  |  |
|  | Marley | -0.47 (-1.31, 0.42) |  |  |  | Marley | 0.01 (-1.59, 1.57) |  |  |
| *Frass Half-Fall ~ Half-Leaf* |  |  |  |  | *Frass Peak ~ Half-Leaf* |  |  |  |  |
| n=77 | Intercept | **130.29 (118.86, 141.94)** | 39.5 | -5.6 | n=77 | Intercept | **128.26 (109.29, 148.32)** | 27.5 | -5.4 |
|  | Half-Leaf | 0.08 (-0.01, 0.17) |  |  |  | Half-Leaf | 0.10 (-0.06, 0.25) |  |  |
|  | FG | **2.24 (1.38, 3.10)** |  |  |  | FG | **3.26 (1.76, 4.80)** |  |  |
|  | Marley | -0.89 (-1.95, 0.18) |  |  |  | Marley | -0.54 (-2.42, 1.28) |  |  |
| *Frass Half-Fall ~ Half-NDVI* |  |  |  |  | *Frass Peak ~ Half-NDVI* |  |  |  |  |
| n=76 | Intercept | **114.43 (98.25, 130.28)** | 45.0 | -0.7 | n=76 | Intercept | **100.60 (73.36, 128.46)** | 34.2 | 0.0 |
|  | Half-NDVI | **0.21 (0.08, 0.35)** |  |  |  | Half-NDVI | **0.33 (0.09, 0.55)** |  |  |
|  | FG | **2.28 (1.49, 3.08)** |  |  |  | FG | **3.37 (1.93, 4.81)** |  |  |
|  | Marley | -0.79 (-1.71, 0.14) |  |  |  | Marley | -0.40 (-2.08, 1.26) |  |  |
| *TC Half-Fall ~ Budburst* |  |  |  |  | *TC Peak ~ Budburst* |  |  |  |  |
| n=77 | Intercept | **125.32 (118.43, 132.29)** | 56.7 | 0.0 | n=77 | Intercept | **107.39 (92.99, 121.75)** | 38.8 | -0.9 |
|  | Budburst | **0.17 (0.11, 0.23)** |  |  |  | Budburst | **0.35 (0.22, 0.48)** |  |  |
|  | FG | **2.15 (1.39, 2.91)** |  |  |  | FG | 1.45 (-0.18, 3.12) |  |  |
|  | Marley | **-1.22 (-2.06, -0.37)** |  |  |  | Marley | **-2.06 (-3.83, -0.26)** |  |  |
| *TC Half-Fall ~ Half-Leaf* |  |  |  |  | *TC Peak ~ Half-Leaf* |  |  |  |  |
| n=77 | Intercept | **128.63 (115.81, 141.15)** | 44.0 | -10.0 | n=77 | Intercept | **117.33 (91.46, 142.81)** | 20.9 | -11.1 |
|  | Half-Leaf | **0.13 (0.03, 0.23)** |  |  |  | Half-Leaf | **0.24 (0.03, 0.45)** |  |  |
|  | FG | **1.98 (1.04, 2.90)** |  |  |  | FG | 1.26 (-0.35, 2.81) |  |  |
|  | Marley | **-1.90 (-3.00, -0.78)** |  |  |  | Marley | **-2.89 (-4.67, -1.09)** |  |  |
| *TC Half-Fall ~ Half-NDVI* |  |  |  |  | *TC Peak ~ Half-NDVI* |  |  |  |  |
| n=76 | Intercept | **107.92 (91.54, 123.67)** | 52.6 | -1.8 | n=76 | Intercept | **64.97 (33.17, 97.43)** | 37.4 | 0.0 |
|  | Half-NDVI | **0.30 (0.17, 0.44)** |  |  |  | Half-NDVI | **0.68 (0.41, 0.94)** |  |  |
|  | FG | **2.06 (1.23, 2.90)** |  |  |  | FG | 1.31 (-0.42, 3.04) |  |  |
|  | Marley | **-1.67 (-2.60, -0.74)** |  |  |  | Marley | **-3.00 (-4.89, -1.01)** |  |  |
| *FWM Half-Fall ~ Budburst* |  |  |  |  | *FWM Peak ~ Budburst* |  |  |  |  |
| n=76 | Intercept | **123.40 (113.91, 133.18)** | 37.8 | 0.0 | n=76 | Intercept | **105.46 (86.65, 124.63)** | 36.0 | 0.0 |
|  | Budburst | **0.20 (0.11, 0.28)** |  |  |  | Budburst | **0.37 (0.20, 0.54)** |  |  |
|  | FG | **1.15 (0.05, 2.25)** |  |  |  | FG | **3.01 (0.87, 5.11)** |  |  |
|  | Marley | **-1.67 (-2.90, -0.41)** |  |  |  | Marley | -2.16 (-4.61, 0.29) |  |  |
| *FWM Half-Fall ~ Half-Leaf* |  |  |  |  | *FWM Peak ~ Half-Leaf* |  |  |  |  |
| n=76 | Intercept | **129.85 (114.25, 145.62)** | 24.6 | -7.8 | n=76 | Intercept | **112.37 (79.66, 143.97)** | 24.5 | -6.6 |
|  | Half-Leaf | **0.13 (0.00, 0.25)** |  |  |  | Half-Leaf | **0.28 (0.03, 0.55)** |  |  |
|  | FG | 1.03 (-0.19, 2.26) |  |  |  | FG | **2.65 (0.20, 5.09)** |  |  |
|  | Marley | **-2.36 (-3.77, -0.93)** |  |  |  | Marley | **-3.70 (-6.62, -0.83)** |  |  |
| *FWM Half-Fall ~ Half-NDVI* |  |  |  |  | *FWM Peak ~ Half-NDVI* |  |  |  |  |
| n=75 | Intercept | **109.92 (86.95, 131.66)** | 30.3 | -2.6 | n=75 | Intercept | **72.74 (30.31, 115.68)** | 31.0 | -0.6 |
|  | Half-NDVI | **0.29 (0.11, 0.48)** |  |  |  | Half-NDVI | **0.62 (0.26, 0.97)** |  |  |
|  | FG | 1.08 (-0.08, 2.26) |  |  |  | FG | **2.98 (0.69, 5.31)** |  |  |
|  | Marley | **-2.14 (-3.45, -0.89)** |  |  |  | Marley | **-3.00 (-5.53, -0.45)** |  |  |

In terms of spatial effects, metrics derived from water trap data (total caterpillar and final instar winter moth half-falls and peaks) tended to be significantly earlier in Marley than Great Wood, though the frass metrics did not significantly deviate between these areas. Instead, ForestGEO oaks were significantly later than Great Wood in all frass models, and for total caterpillar half-fall and final instar winter moth peak models. Despite these area effects, the variation in caterpillar metrics explained by models using either budburst or half-NDVI as the oak tree phenology metric was very similar for frass half-falls and peaks, and total caterpillar half-falls and peaks. However, budburst explained slightly more variation than half-NDVI in the final instar winter moth models. Half-leaf always explained the least variation. For example, in frass half-fall models, budburst explained 47.7% of the variance, half-NDVI 45.0%, and half-leaf 39.5%. This pattern was mirrored in the ELPD-LOO scores (**Table 1**).

### Relationships between individual oak tree or caterpillar phenology and herbivory rates

Herbivory, measured as the mean percentage of leaf area consumed across ten leaves for 72 oaks, showed weak to moderate negative correlations with tree and caterpillar phenology metrics, indicating that trees with later phenology had lower herbivory levels. Significant associations were observed with half-leaf date (r=-0.40, p<0.001, the strongest correlation), half-NDVI (r=-0.26, p<0.05), frass half-fall (r=-0.24, p<0.05) and frass peak dates (r=-0.28, p<0.05; all n=72). Models controlling for spatial and tree identity effects showed that half-leaf explained the most variation in herbivory compared to other phenological metrics, with a conditional *R*^2^ value of 33.3%, followed by half-NDVI (*R*^2^=26.6%), frass half-fall (*R*^2^=26.1%) and frass peak (*R*^2^=24.9%). Only models with half-leaf, frass half-fall or peak had significant negative effects on herbivory after controlling for sampling area and branch-level variation, with frass half-fall having the strongest effect (**Table 2**; **SI Figure 6**). The model for half-NDVI was marginally non-significant. The pattern is further supported by the ELPD-LOO scores, with half-leaf performing the best. Additionally, eight of the nine models (all except half-leaf) showed significant effects of sampling area, with oaks in ForestGEO and Marley having significantly lower herbivory levels compared to those in Great Wood.

**Table 2.** Outputs of Bayesian Beta mixed models for the effect of tree and caterpillar phenology metrics on herbivory for 72 oak trees in Wytham Woods. Note that estimates are on the logit scale. All models included one phenology metric and sampling area as fixed effects and controlled for the relative canopy level (lower, middle, upper) that the sampled leaves came from. Intercept is taken as area = Great Wood. 95%CIs are 95% credible intervals around the estimate and effects in bold are significant where CIs do not overlap zero. *R*^2^ is the amount of variation explained by the model (conditional i.e. includes the random effect of canopy level). Δelpd is the difference in ELPD-LOO between the best fitting model (set to 0) and all others. HF = half-fall, TC = total caterpillars, FWM = final instar winter moth caterpillars.

|  | Estimate (95%CI) | R<sup>2</sup> (%) | Δelpd |
| --- | --- | --- | --- |
| *Herbivory ~ Budburst* |  |  |  |
| n=72 | Intercept -0.18 (-4.21, 3.40) | 21.7 | -3.7 |
|  | Budburst -0.02 (-0.05, 0.02) |  |  |
|  | FG **-0.55 (-0.97, -0.11)** |  |  |
|  | Marley **-0.69 (-1.20, -0.19)** |  |  |
| *Herbivory ~ Half-Leaf* |  |  |  |
| n=72 | Intercept **6.27 (0.54, 11.91)** | 33.3 | 0.0 |
|  | Half-Leaf **-0.07 (-0.11, -0.02)** |  |  |
|  | FG -0.43 (-0.87, 0.02) |  |  |
|  | Marley -0.35 (-0.91, 0.21) |  |  |
| *Herbivory ~ Half-NDVI* |  |  |  |
| n=71 | Intercept 5.42 (-3.06, 13.55) | 26.6 | -4.7 |
|  | Half-NDVI -0.06 (-0.13, 0.01) |  |  |
|  | FG **-0.55 (-0.99, -0.11)** |  |  |
|  | Marley **-0.66 (-1.18, -0.15)** |  |  |
| *Herbivory ~ Frass Half-Fall* |  |  |  |
| n=72 | Intercept **17.64 (0.69, 34.96)** | 26.1 | -0.05 |
|  | Frass HF **-0.14 (-0.26, -0.02)** |  |  |
|  | FG -0.20 (-0.75, 0.33) |  |  |
|  | Marley **-0.73 (-1.45, -0.24)** |  |  |
| *Herbivory ~ Total Caterpillar Half-Fall* |  |  |  |
| n=72 | Intercept 6.70 (-7.87, 22.58) | 21.5 | -3.3 |
|  | TC HF -0.06 (-0.17, 0.04) |  |  |
|  | FG -0.42 (-0.93, 0.10) |  |  |
|  | Marley **-0.73 (-1.45, -0.24)** |  |  |
| *Herbivory ~ Final Instar Winter Moth Half-Fall* |  |  |  |
| n=71 | Intercept -2.17 (-13.11, 7.92) | 21.0 | -5.4 |
|  | FWM HF 0.00 (-0.07, 0.08) |  |  |
|  | FG **-0.57 (-1.00, -0.12)** |  |  |
|  | Marley **-0.70 (-1.28, -0.19)** |  |  |
| *Herbivory ~ Frass Peak* |  |  |  |
| n=72 | Intercept 8.98 (-0.61, 19.58) | 24.9 | -0.9 |
|  | Frass Peak **-0.08 (-0.15, -0.01)** |  |  |
|  | FG -0.29 (-0.80, 0.20) |  |  |
|  | Marley **-0.67 (-1.18, -0.18)** |  |  |
| *Herbivory ~ Total Caterpillar Peak* |  |  |  |
| n=72 | Intercept 2.41 (-4.83, 10.16) | 21.3 | -4.1 |
|  | TC Peak -0.03 (-0.08, 0.02) |  |  |
|  | FG **-0.51 (-0.98, -0.05)** |  |  |
|  | Marley **-0.70 (-1.23, -0.22)** |  |  |
| *Herbivory ~ Final Instar Winter Moth Peak* |  |  |  |
| n=71 | Intercept -1.26 (-6.90, 4.00) | 20.8 | -6.3 |
|  | FWM Peak -0.00 (-0.04, 0.03) |  |  |
|  | FG **-0.55 (-1.02, -0.10)** |  |  |
|  | Marley **-0.70 (-1.25, -0.20)** |  |  |

## Discussion

Different methods of measuring individual phenology may or may not adequately capture spatiotemporal variation in phenological relationships. We derived three metrics of tree phenology and six metrics for the timing of phytophagous larvae from five field methods performed on 170 trees in a deciduous woodland and assessed relationships between metric values, both within and between trophic levels, and tested the extent to which they can explain herbivory as the outcome of trophic interactions.

The six common tree species sampled in this study differed in absolute dates for nine metrics and the orders they reached these events (**Figure 4**; **SI Table 2**). The order that species reached budburst was identical to that which Cole & Sheldon (2017) found in a larger sample of trees (total n=825) at the same study site ten years previously. They found the spread in species budburst means to be 20 days in 2013 versus 42 days in 2014, whilst they are intermediate in our study with 32 days between hazel and ash budburst (Cole & Sheldon, 2017). Hence, this comparison suggests our data were collected in a somewhat typical year for phenological variation. Species explains the most variation in budburst compared to other metrics (*R*^2^=73.1%), and this specieslevel variation in tree phenology can be attributed to species-specific responses to environmental cues (Vitasse et al., 2009; Roberts et al., 2015). Half-leaf and half-NDVI exhibited narrower ranges of variation, suggesting that budburst is highly species-dependent whereas subsequent phenological events may be more synchronised across species. However, the small amount of variation explained by species for half-leaf (*R*^2^=19.1%) could be due to the influence of non-focal tree species within the image frame. This analysis further emphasises the relevance of interspecific phenological variation promoting complexity in the woodland phenological landscape.

Different timing of our phenological metrics was expected given that they measure different stages in a dynamic process. We found that half-NDVI was consistently later than budburst across species, and half-leaf later again (**Figure 4**; **SI Table 2**). Budburst is when small leaves first start to emerge, which would not be picked up as 50% of maximum NDVI until they have emerged and grown larger, hence half-NDVI day occurs later. Other studies validating aerial imagery with ground observations have also found that NDVI values lag behind budburst or start of growing season metrics (Soudani et al., 2008; Berra et al., 2019). The strong correlation between budburst and half-NDVI in oak and sycamore, and to a lesser extent in ash and birch (**Figure 5d & e, a & b**), suggests that oak leaf development follows a relatively predictable trajectory. However the relationship is variably correlated, which suggests these events may decouple i.e. leaf emergence rates differ among individuals within species, or even between different parts of the same canopy (e.g. in terms of a structural order where leaves may emerge earlier or faster in the lower canopy to avoid shading from those emerging at the top). The same could be argued for half-leaf being on average later than budburst and half-NDVI but not correlated with either of them for ash, birch, oak and sycamore trees, however we believe the lack of correlation is due mostly to methodological reasons (see below).

While half-dates are widely used in phenological research as a standardised way to compare timing, their application can have limitations across different processes. For example, budburst occurs earlier in spring than half-NDVI or half-leaf, making it more susceptible to stochastic weather events such as cold spells which can pause bud development. This results in deviations from the sigmoidal pattern assumed by half-date extraction models (as illustrated by the data for an individual oak tree in **Figure 3**). In contrast, the NDVI and LAI measurements begin increasing later in the season, after the most variable early-spring weather events, and tend to follow smoother sigmoid trajectories. Such deviations may increase uncertainty in calculating budburst dates and contribute to their variability compared to the later season tree phenology metrics like half-NDVI and half-leaf (**SI Table 2**). Although budburst timing is likely to be the event under selection for synchrony with caterpillar hatching, the accuracy with which it can be quantified using bud observations alone may limit precise comparisons with metrics that reflect cumulative consumer activity. Future work may benefit from exploring alternative or complementary parameters of phenological distributions (Macphie et al., 2024).

At the caterpillar level, phenology metrics were much more consistent across host tree species, indicating caterpillar timing may be more influenced by other factors such as local environmental/microclimatic conditions and individual tree characteristics. There are also small differences between the mean dates of metrics within tree species, with final instar winter moth metrics being 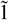 day later than total caterpillar, which are 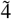 days later than frass metrics, and this is true for both half-fall and peak dates. Final instar winter moth half-fall had the widest credible intervals around the date estimates in non-oaks (**SI Table 2**), as their lower abundance affects the fitting of the cumulative curves needed to estimate half-fall date and reduced the sample sizes as some trees collected none at all. There was strong correlation between all caterpillar metrics in individual oak trees, but weaker relationships in non-oaks (**Figure 5**). Water traps will not capture the total composition of larvae feeding in the canopy, as only late instar larvae of species descending to pupate, or individuals that fall off leaves, will be collected; this bias explains why total caterpillar and final instar winter moth dates occurred later. Frass fall, whilst lacking caterpillar species information, is likely the more accurate representation of all larvae in the canopy at any given time – especially if temperature-corrected to estimate biomass (Tinbergen & Dietz, 1994; Hinks et al., 2015; Tinbergen et al., 2024). Given also that frass is the product of herbivory, this explains why frass was significantly related to oak herbivory levels whilst water trap metrics were not (**Table 2**; **SI Figure 6**). However, there is a logistical drawback to using frass traps, which is that heavy rain can dissolve collected pellets or prevent them from falling off leaves into traps. This means it may only be possible to collect adequate frass time series during unusually dry springs, which we were fortunate to have at our site in 2023.

Between trophic levels, oaks showed the strongest correlations between tree and caterpillar phenology (**Figure 6**), which aligns with the relatively high caterpillar abundance typically observed on oaks compared to the other co-occurring tree species we sampled (mean peak abundance of total caterpillars under oaks was 31.50, versus: ash = 3.42, beech = 5.24, birch = 4.64, hazel = 8.88, and sycamore = 17.54; see also Shutt et al., 2019; Macphie et al., 2024). We found that oak budburst showed slightly stronger correlation with final instar winter moth half-fall than with frass peak dates (BB–FWMHF r=0.50, p<0.001; n=75; BB–FP r=0.39, p<0.001; n=76), consistent with Hinks et al. (2015). The lack of strong correlations between tree and caterpillar phenology for other host tree species might imply that these relationships are less tightly coupled in those species. However, it may also result from a lower sample size in this study, as well as lower abundance of insect herbivores on these host trees affecting the precision of phenological metric estimates. Within oaks, budburst and half-NDVI date both have significant positive effects and explain similar proportions of variation in frass and total caterpillar phenology, although half-NDVI always has the stronger effect (**Table 1**). For final instar winter moth metrics, half-NDVI again has the stronger effect but explains slightly less variation than budburst. Statistically, budburst or half-NDVI would be a good method of measuring tree phenology in the context of tree-invertebrate interactions, although logistically there are pros and cons to each. Bud scoring may be subjective but training observers and controlling for observer effects in the extraction models can mitigate that. Sample sizes are limited by the logistics of ground-scoring but calculating budburst date involves virtually no data processing and fitting the extraction curves was less noisy than the two technological methods. Drone imagery requires substantial processing steps to derive vegetation index data, considerable expense, and is dependent on suitable weather conditions for flying, but could be applied to much larger sample sizes (whether through manual crown delimitation or future automatic segmentation; Berra et al., 2019).

The weaker effects of half-leaf and the fact that it explains the least variation in all six caterpillar metrics, (**Table 1**), indicates that despite offering less subjective quantification of leaf emergence at ground level, it is the least suitable method of monitoring tree phenology in the context of tree-invertebrate interactions. The reason for this could be partly biological, in that the appearance of fresh leaves with good nutritional quality at budburst is crucial for the growth and survival of newly hatched larvae, so budburst is the event under stronger selection for synchrony than later leaf expansion (Tikkanen & Julkunen-Tiitto, 2003; van Asch & Visser, 2007), but half-NDVI is still a significant predictor of caterpillar metrics because of its high correlation with oak budburst (**Figures 5d & 6d**). The unsuitability of half-leaf to predict caterpillar phenology may also be partly methodological, in that raw LAI values at the time of budburst vary depending on individual tree attributes (e.g. species, canopy size, branch density), so there is no threshold of actual or standardised LAI across trees that would match with budburst date. Secondly, the nature of hemispherical photography means that non-focal individuals (both trees and understorey) in the edges of the frame can skew the overall image’s LAI values, hence the lack of correlation to bud score observations of individual trees or segmentation of individual crowns in aerial images. Half-leaf was only correlated with budburst in hazel trees because, as understorey trees, their canopies are closer to the ground camera and in the foreground of the hemispherical photos, resulting in less non-focal vegetation in frame, therefore correlating better with direct bud observations compared to higher canopy trees (**Figure 5c**). Segmentation of the focal tree in hemispherical images might improve this (e.g. leaf metrics based on visual scoring corresponded well to non-hemispherical PhenoCam images with defined “regions of interest”, Xie et al., 2018), as could accounting for uncertainty (Brown et al., 2023), but hemispheR currently lacks the ability to perform on segmented hemispherical images.

Furthermore, despite half-leaf exhibiting no significant correlation with other tree or caterpillar metrics, it appeared to only very slightly better explain variation in oak herbivory (**Table 2**; **SI Figure 6**). This may be because LAI is affected by leaf clumping, so the density of this food resource may affect herbivory levels. Even where metric effects are non-significant, all models indicate that trees with later phenology have lower herbivory, aligning with the expectations of Crawley & Akhteruzzaman (1988) although they did not find the expected significant correlation in their population. All models left substantial variation in herbivory unexplained (**Table 2**). Other factors such as palatability, light environment and larval abundance/density, or the actual number of days mismatch/degree of (a)synchrony between tree and caterpillar phenology, rather than direct phenology metrics, likely contribute to this variation in herbivory among trees (Tikkanen & Julkunen-Tiitto, 2003; Barber & Marquis, 2011; Crawley & Akhteruzzaman, 1988).

There is some evidence to suggest spatial variation in phenology, as well as the impact of herbivory, within the study site here (**Tables 1 & 2**; **SI Figures 5 & 6**). Such spatial variation in phenology and herbivory may result from differential woodland structures, tree species compositions, or microclimatic effects for example that affect larval abundance/density (Heinecke et al., 2024). Combined with the species-specific and individual-level findings, this suggests the phenological landscape within this single study system is complex and varies at fine spatial scales. This further reinforces the need to go beyond population means and account for individual variation when predicting the consequences of phenological (mis)match under changing climates.

One limitation of this study is that data were collected over a single year, and phenological relationships may vary depending on inter-annual temperature trends, hence our findings may not generalise across years with distinctly different weather conditions. Repeated multi-method datasets across multiple years would be required to assess the temporal consistency of these relationships under varying environmental conditions. We also did not directly measure caterpillar abundance through branch sampling due to logistical constraints (although Visser et al. (2006) found that biomass estimated from branches and from frass traps was highly correlated (*F*_1,13_=19.35, p<0.001)). Whilst logistically easier to deploy at scale, water and frass traps only measure proxies of the caterpillar abundance and community composition that exists in tree canopies. Frass traps better reflect canopy herbivory, but they cannot differentiate species, while water traps may be biased towards later instar individuals of the subset of species which fall to the ground to pupate, underestimating the true larval canopy abundance. Future work comparing branch sampling against these indirect ground measures will clarify their validity.

Among the methods we applied, our findings suggest that bud scoring is a robust and low-cost option for monitoring tree phenology and is well-correlated with caterpillar timing and herbivory but has sample size limitations. NDVI derived from drone imagery, while more logistically demanding, scales up sampling well and captures the development of canopy green-up that also correlates strongly with caterpillar metrics, particularly in oaks. In contrast, LAI measured by hemispherical photography appears less suitable for capturing individual tree phenology and synchrony with primary consumers, both biologically and methodologically. For caterpillar phenology, frass traps provide the more comprehensive measure of their indirect biomass, as long as the site does not have high rainfall, while water traps can offer useful species-level data but may underestimate their true abundance in the canopy. The choice of method should therefore be guided by which type of data best answers a given study’s ecological question and its logistical constraints, although integrating multiple methods could help balance trade-offs between accuracy, scalability and ecological relevance.

In conclusion, the field methods used to monitor the phenology of individual trees and their phytophagous larvae communities can yield different estimated values of timing within trophic levels. In oaks, which have been a particular focus of much work, despite consistent qualitative relationships in phenology between trophic levels and with herbivory, their strength and quantitative outputs exhibit some variation depending on the phenological metrics chosen. Relationships are generally weaker in non-oak tree species. As each method captures slightly different aspects of phenological processes, selecting the appropriate approach should be guided by which type of data best answers the ecological question being explored and any logistical constraints of the study and site. Although, integrating multiple methods could help balance trade-offs between accuracy, scalability and ecological relevance. This analysis highlights the importance of methodological considerations in capturing phenological dynamics of trophic interactions within woodland ecosystems.

## Data Accessibility Statement

The data and analysis scripts that support the findings of this study are available on Zenodo at DOI: https://doi.org/10.5281/zenodo.14721158

## Author Contributions

LMM, EFC and BCS contributed to the ideas of this study. LMM collected data and conducted the statistical analyses with inputs on drone data methodology from SJC. LMM wrote the manuscript with EFC, BCS and SJC providing feedback. All authors approved the final manuscript. LMM: Conceptualization; writing - original draft; formal analysis; writing - review and editing. SJC: Resources; software; writing - review and editing. EFC: Conceptualization; writing - review and editing. BCS: Conceptualization; writing - review and editing.

## ACKNOWLEDGEMENTS

We thank Gwen Lewis, Freya Coursey, Leo Fordham, Alex Rosenfeld, Chenjing Huang, Josh Hill and Sienna Rattigan for their help with the field data collection, and the PhenoScale group (Department of Biology, University of Oxford) for their comments during the development of this work. This work was funded by a UKRI Frontiers grant EP/X024520/1 to BCS.

