## Supplementary Information for "Quantifying phenology in the deciduous tree and phytophagous insect system: a methodological comparison"

| Field Method | Start Date | End Date | Number of Sampling Days |
|---|---|---|---|
| Multispectral drone imaging | 21st March | 9th June | 21-23 |
| Bud scoring observations | 25th March | 25th May | 20-21 |
| Hemispherical canopy photography | 25th March | 8th June | 16-22 |
| Water traps | 3rd May | 20th June | 14 |
| Frass traps | 9th May | 15th June | 11 |

**Supplementary Table 1.** Collection dates for each of the five field methods measuring the tree and caterpillar phenology of 170 trees in Wytham Woods, Oxfordshire, during spring 2023. Variation in the number of sampling days depends on flight/ground sampling area.

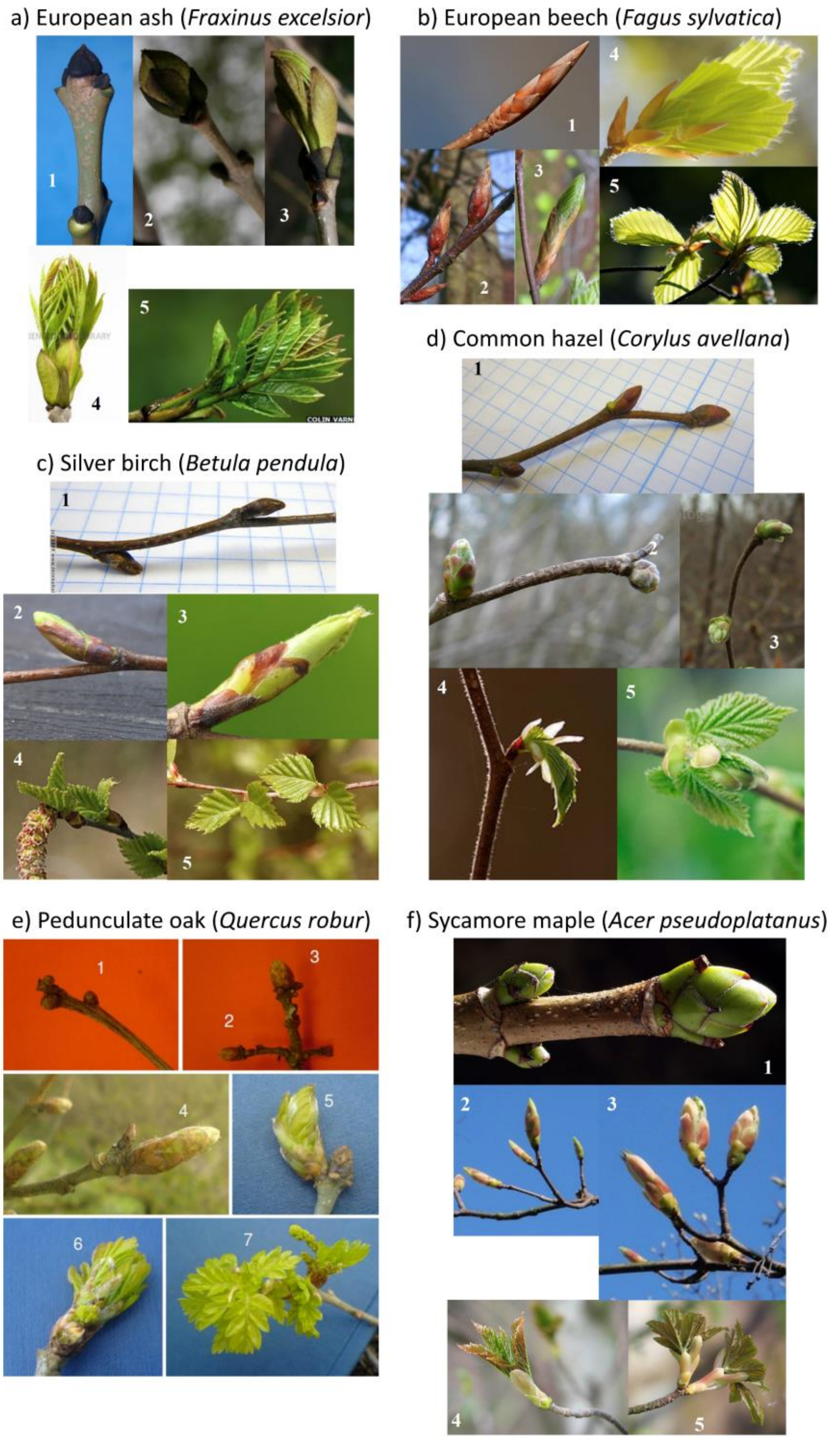

**Supplementary Figure 1.** Photographic reference keys used for scoring bud development, from 1 (no or minimal bud swelling), to 5 (for non-oaks; **a-d & f**) or 7 (for oaks; **e**) denoting full leaf. These are the same keys used in Cole & Sheldon (2017). Images taken from Google Images.

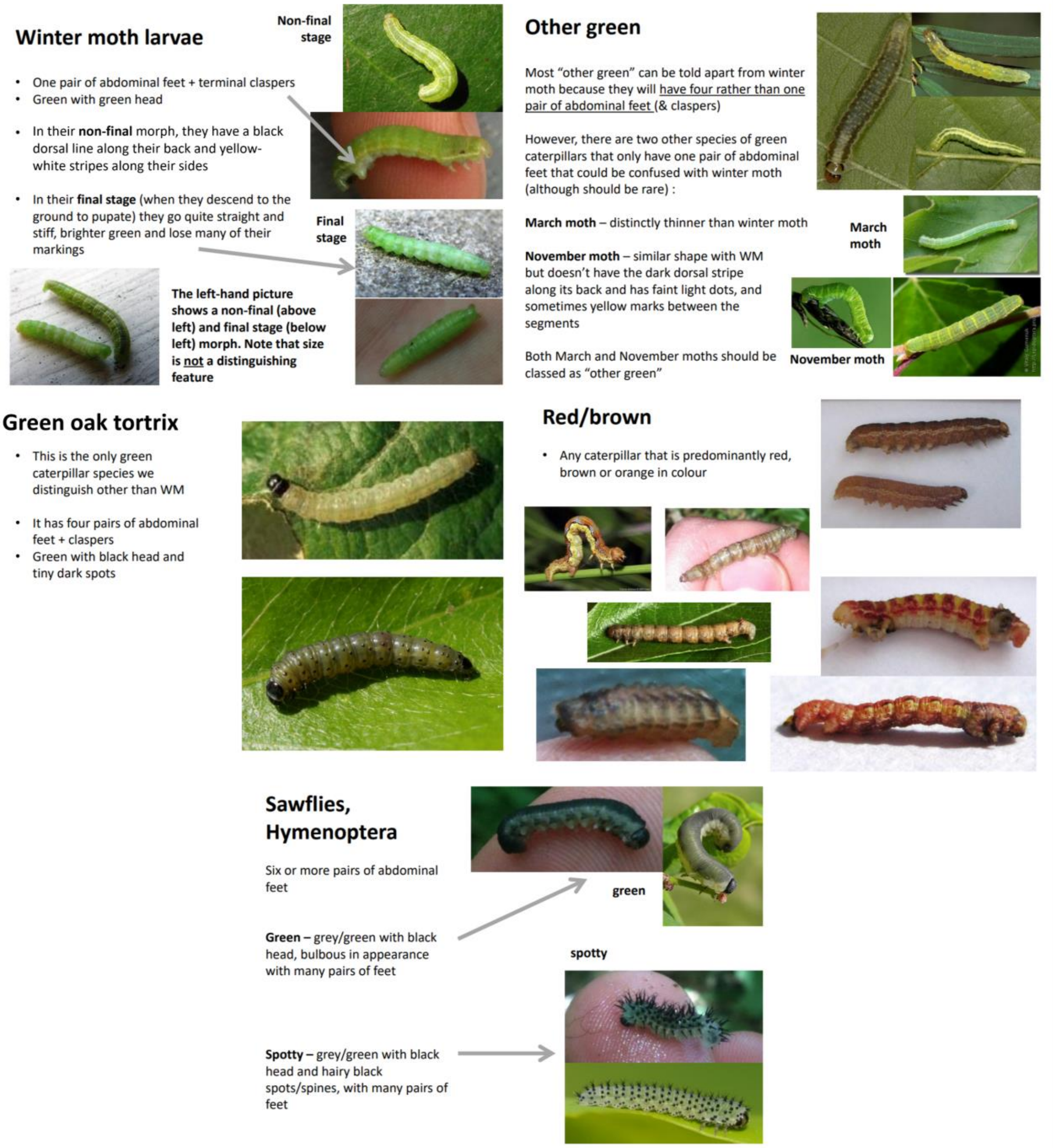

**Supplementary Figure 2.** Photographic reference keys used for identifying caterpillar groups. Images taken from Google Images.

| Species | Metric | Mean Day (±sd) | Mean CI Width (±sd) | n | Rank |
|---|---|---|---|---|---|
| **Ash** | Budburst | 123.76 (3.34) | 5.11 (1.08) | 23 | 6 |
| | Half-Leaf | 131.96 (4.05) | 8.56 (1.54) | 23 | 6 |
| | Half-NDVI | 126.21 (2.87) | 7.64 (2.13) | 22 | 5 |
| | Frass Half-Fall | 143.23 (2.23) | 3.96 (0.74) | 23 | 6 |
| | TC Half-Fall | 146.78 (3.37) | 7.91 (3.13) | 22 | 5 |
| | FWM Half-Fall | 149.10 (3.14) | 14.18 (2.91) | 12 | 6 |
| | Frass Peak | 143.14 (3.37) | | 23 | 4 |
| | TC Peak | 147.97 (5.08) | | 22 | 4 |
| | FWM Peak | 151.20 (5.03) | | 12 | 6 |
| **Beech** | Budburst | 110.79 (4.52) | 10.67 (3.34) | 5 | 4 |
| | Half-Leaf | 125.06 (1.24) | 6.52 (1.98) | 5 | 2 |
| | Half-NDVI | 121.37 (1.20) | 4.85 (0.57) | 5 | 4 |
| | Frass Half-Fall | 142.75 (1.25) | 3.81 (0.31) | 5 | 4 |
| | TC Half-Fall | 143.07 (4.24) | 6.01 (2.26) | 5 | 1 |
| | FWM Half-Fall | 143.97 (3.05) | 12.12 (1.05) | 4 | 1 |
| | Frass Peak | 143.95 (2.89) | | 5 | 6 |
| | TC Peak | 143.27 (3.91) | | 5 | 1 |
| | FWM Peak | 145.55 (3.30) | | 4 | 1 |
| **Birch** | Budburst | 100.33 (3.95) | 6.84 (2.61) | 17 | 2 |
| | Half-Leaf | 125.93 (5.07) | 8.87 (1.75) | 17 | 3 |
| | Half-NDVI | 115.25 (3.20) | 8.64 (2.28) | 16 | 2 |
| | Frass Half-Fall | 141.05 (1.83) | 3.58 (0.69) | 17 | 2 |
| | TC Half-Fall | 144.09 (4.11) | 5.88 (0.10) | 16 | 2 |
| | FWM Half-Fall | 146.31 (2.39) | 8.97 (2.90) | 14 | 3 |
| | Frass Peak | 140.41 (1.40) | | 17 | 1 |
| | TC Peak | 147.42 (5.34) | | 16 | 3 |
| | FWM Peak | 149.63 (4.60) | | 14 | 4 |
| **Hazel** | Budburst | 91.86 (5.36) | 11.52 (2.63) | 26 (CI n=6)* | 1 |
| | Half-Leaf | 127.28 (3.72) | 7.45 (1.34) | 30 | 5 |
| | Half-NDVI | NA | NA | NA | NA |
| | Frass Half-Fall | 141.58 (1.93) | 3.25 (0.63) | 30 | 3 |
| | TC Half-Fall | 146.69 (3.22) | 7.48 (2.23) | 30 | 4 |
| | FWM Half-Fall | 148.33 (2.43) | 11.15 (2.67) | 24 | 4 |
| | Frass Peak | 141.59 (1.96) | | 30 | 3 |
| | TC Peak | 149.76 (5.62) | | 30 | 5 |
| | FWM Peak | 150.37 (4.17) | | 24 | 5 |
| **Oak** | Budburst | 111.88 (5.50) | 11.07 (5.28) | 77 | 5 |
| | Half-Leaf | 125.94 (4.38) | 6.90 (1.42) | 77 | 4 |
| | Half-NDVI | 121.27 (2.82) | 5.88 (1.33) | 76 | 3 |
| | Frass Half-Fall | 140.73 (2.04) | 1.68 (0.35) | 77 | 1 |
| | TC Half-Fall | 145.03 (2.21) | 2.20 (0.39) | 77 | 3 |
| | FWM Half-Fall | 145.42 (2.56) | 5.84 (1.30) | 76 | 2 |
| | Frass Peak | 141.24 (3.27) | | 77 | 2 |
| | TC Peak | 146.83 (3.89) | | 77 | 2 |
| | FWM Peak | 147.70 (4.98) | | 76 | 2 |
| **Sycamore** | Budburst | 102.86 (6.85) | 8.20 (2.75) | 18 (CI n=17)* | 3 |
| | Half-Leaf | 123.56 (3.11) | 6.69 (0.70) | 18 | 1 |
| | Half-NDVI | 114.15 (4.16) | 6.52 (1.45) | 15 | 1 |
| | Frass Half-Fall | 142.83 (2.28) | 2.51 (0.33) | 18 | 5 |
| | TC Half-Fall | 147.61 (2.14) | 3.91 (0.70) | 17 | 6 |
| | FWM Half-Fall | 148.78 (2.95) | 11.09 (2.39) | 17 | 5 |
| | Frass Peak | 143.18 (2.51) | | 18 | 5 |
| | TC Peak | 150.55 (5.04) | | 17 | 6 |
| | FWM Peak | 149.30 (5.01) | | 17 | 3 |

**Supplementary Table 2.** Means and standard deviations of day values for nine phenological metrics for each tree species across 170 trees sampled in Wytham Woods. Rank denotes the order in which species reach their mean date for each metric, 1 being earliest, 6 being latest (5 for half-NDVI as hazel does not have NDVI data). Reduction in sample size for hazel budburst is due to four trees having curves that would cross y=0.5 earlier than the start of sampling, likewise for the lower sample size in hazel and sycamore budburst CIs as the lower bounds were outside of day range. Lower sample sizes in NDVI occur as we were unable to delineate a minority of tree crowns, whilst lower sample sizes in caterpillar metrics occur when water traps for certain trees never contained any caterpillars or specifically final-instar winter moths.

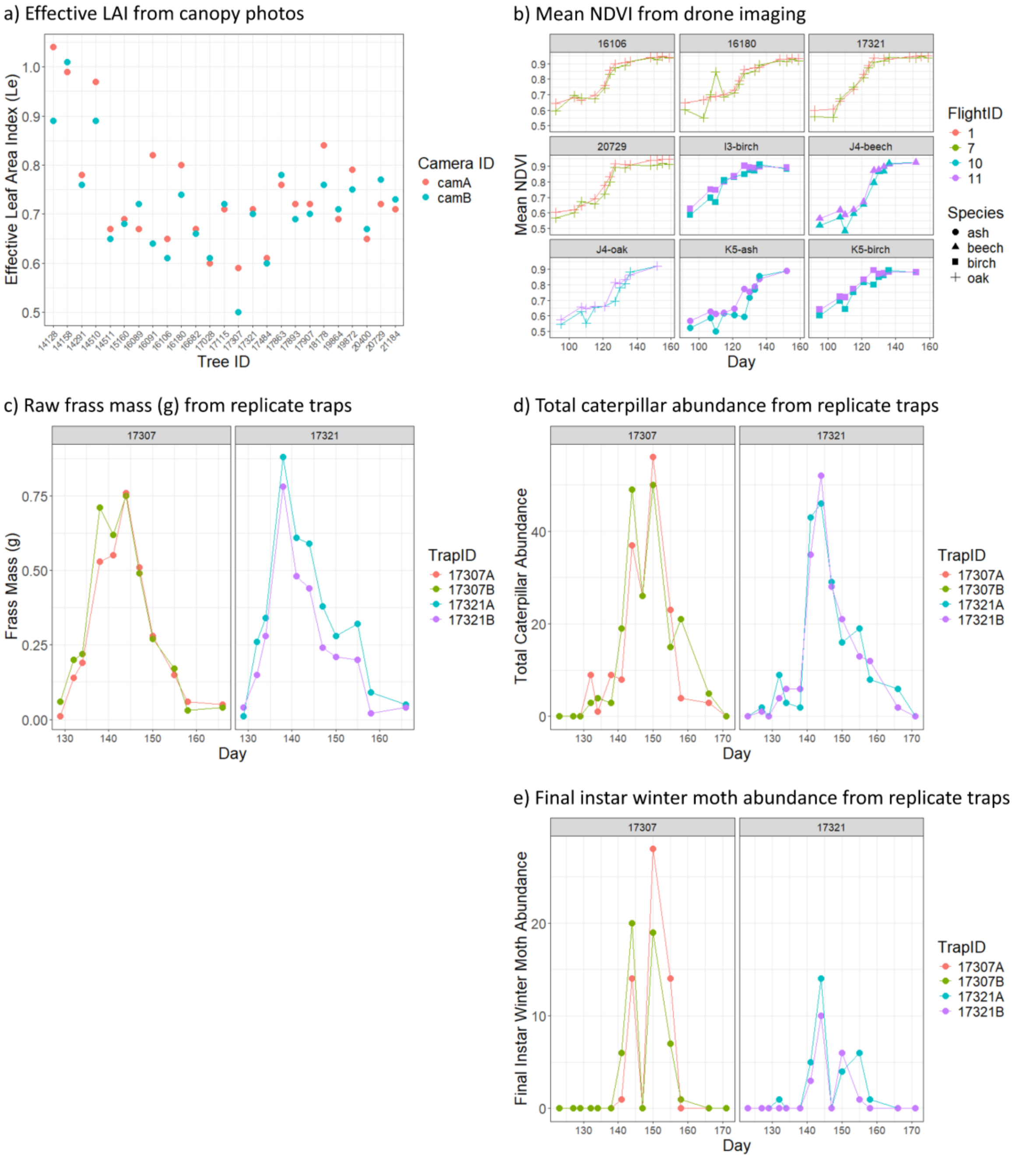

**Supplementary Figure 3.** Visualisations of repeated measurements made to assess repeatability of four out of the five field methods. **a)** The 25 ForestGEO oaks were imaged by two identical camera setups, one immediately after the other, on 28th March 2023. **b)** Nine trees were within overlapping flights so two sets of mean NDVI values existed for when the two flights occurred on the same day. **c)** Two oak trees had two frass traps beneath them across the whole season, and similarly the same two trees also had two replicate water traps to give two sets of **d)** total caterpillar and **e)** final instar winter moth counts.

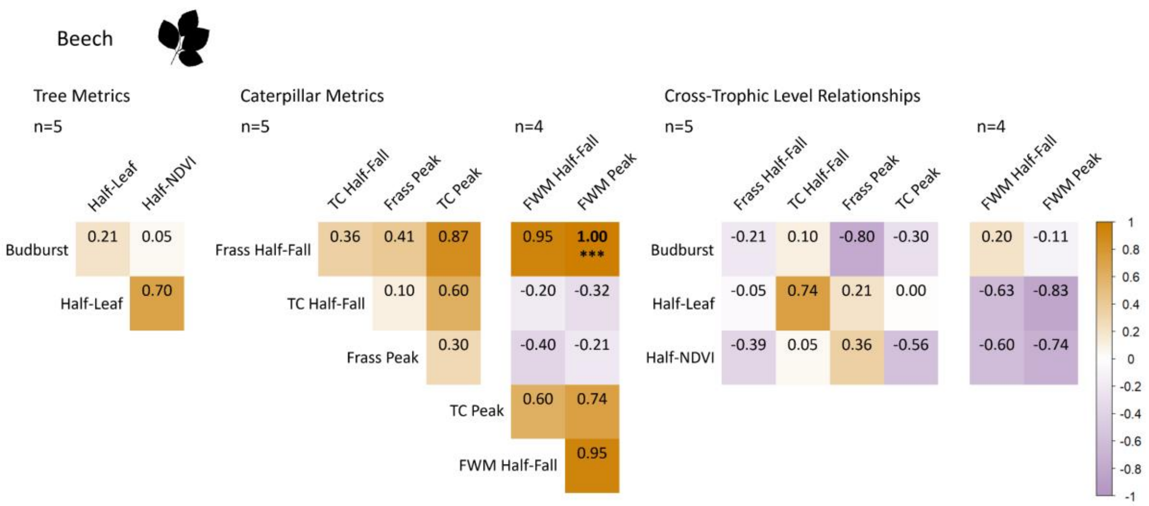

**Supplementary Figure 4.** Correlation matrix between the nine phenological metrics for beech trees. Values are Spearman correlation coefficients; those in bold are significant at the $p<0.05$ threshold and asterisks denote significance thresholds: * = $p<0.05$, ** = $p<0.01$, *** = $p<0.001$. Differing sample sizes between occur as one tree did not collect any final-instar winter moths.

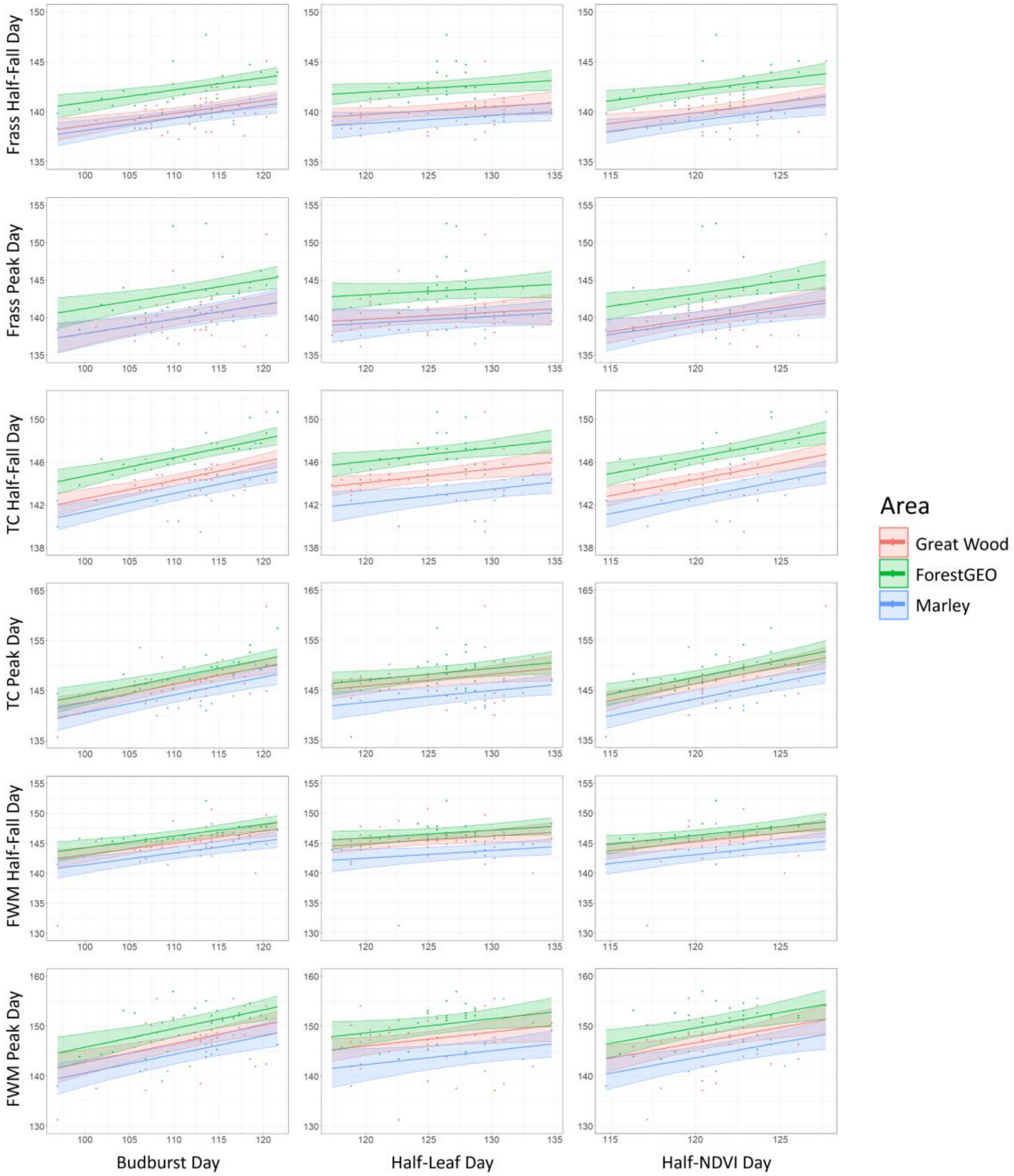

**Supplementary Figure 5.** Conditional effects plots of models quantifying relationships between three tree phenology metrics (columns) and six caterpillar phenology metrics (rows) for 77 oak trees in Wytham Woods, across sampling areas. Shaded areas are 95% credible intervals. Note that x-axis scales are consistent within tree metrics and y-axis scales are consistent within caterpillar metrics. Coefficients, $R^2$ and statistical significance of these relationships are presented in **Table 1**. TC = total caterpillars, FWM = final instar winter moths.

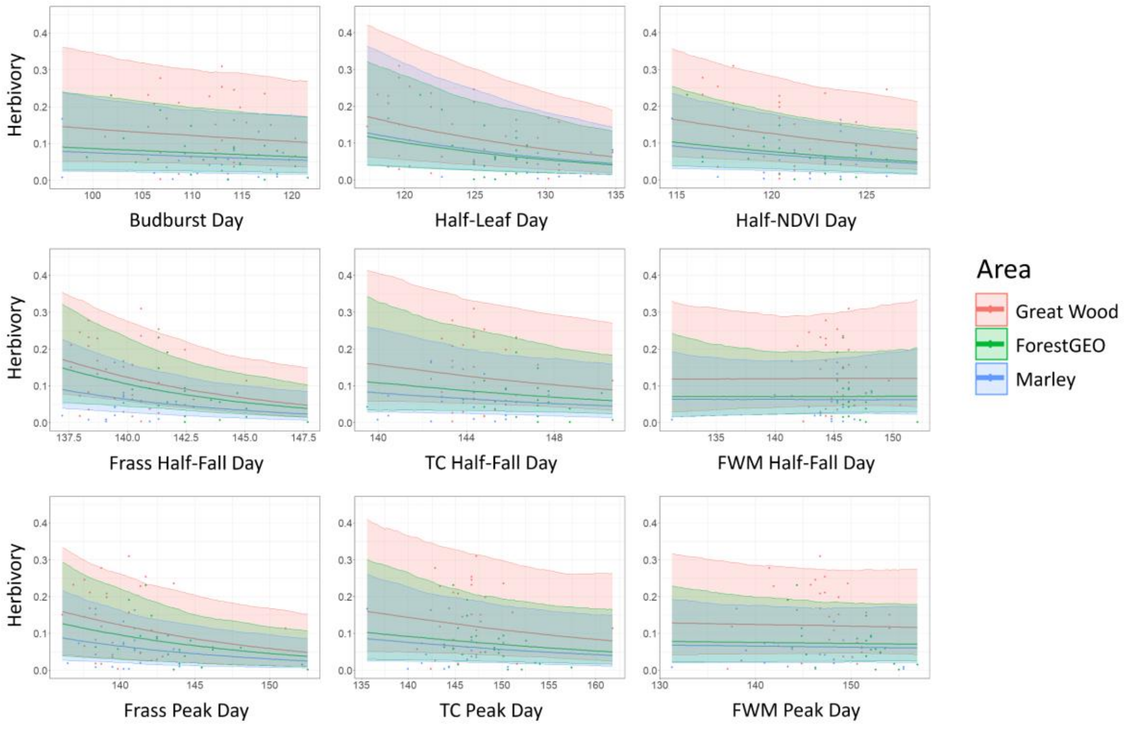

**Supplementary Figure 6.** Conditional effects plots of models quantifying relationships between herbivory and nine phenology metrics of 72 oak trees in Wytham Woods, across sampling areas. Shaded areas are 95% credible intervals. Note that the y-axis scale is consistent between plots but the x-axis scale changes between metrics. Coefficients, $R^2$ and statistical significance of these relationships are presented in **Table 2**. TC = total caterpillars, FWM = final instar winter moths.
